# WNK kinases sense molecular crowding and rescue cell volume via phase separation

**DOI:** 10.1101/2022.01.10.475707

**Authors:** Cary R. Boyd-Shiwarski, Daniel J. Shiwarski, Shawn E. Griffiths, Rebecca T. Beacham, Logan Norrell, Daryl E. Morrison, Jun Wang, Jacob Mann, William Tennant, Eric N. Anderson, Jonathan Franks, Michael Calderon, Kelly A. Connolly, Claire J. Weaver, Claire C. Weckerly, Udai Bhan Pandey, Christopher J. Donnelly, Dandan Sun, Aylin R. Rodan, Arohan R. Subramanya

## Abstract

When challenged by hypertonicity, dehydrated cells must defend their volume to survive. This process requires the phosphorylation-dependent regulation of SLC12 cation chloride transporters by WNK kinases, but how these kinases are activated by cell shrinkage remains unknown. Within seconds of cell exposure to hypertonicity, WNK1 concentrates into membraneless droplets, initiating a phosphorylation-dependent signal that drives net ion influx via the SLC12 cotransporters to rescue volume. The formation of WNK1 condensates is driven by its intrinsically disordered C-terminus, whose evolutionarily conserved signatures are necessary for efficient phase separation and volume recovery. This disorder-encoded phase behavior occurs within physiological constraints and is activated in vivo by molecular crowding rather than changes in cell size. This allows WNK1 to bypass a strengthened ionic milieu that favors kinase inactivity and reclaim cell volume through condensate-mediated signal amplification. Thus, WNK kinases are physiological crowding sensors that phase separate to coordinate a cell volume rescue response.

## Introduction

Eukaryotic cells are constantly subjected to environmental challenges that threaten their fluid volume. During hypertonic stress, cells adapt to volume contraction within seconds by orchestrating a system of ion transporters, channels, and pumps that drive net solute influx, resulting in the obligate reclamation of water (Delpire and Gagnon, 2018). This rapid response, termed regulatory volume increase (RVI), provides a stopgap that gives cells time to synthesize osmolytes and initiate transcriptional programs essential for long-term survival (Burg, 1995). RVI is mediated by specific members of the SLC12 family of electroneutral cation chloride cotransporters (Delpire and Gagnon, 2018). Hypertonic stress activates ion influx via the Na-K- 2Cl cotransporter NKCC1; simultaneously, K-Cl cotransporters (KCCs) are inhibited, blocking ion efflux. This results in the net movement of sodium, potassium, and chloride ions into the cell, which stimulates volume recovery. These reciprocal changes in NKCC1/KCC activity are mediated by serine-threonine phosphorylation of the cotransporters via a common signaling cascade that requires With-No-Lysine (WNK) kinases (de Los Heros et al., 2018). Hyperosmotic stress strongly activates the WNKs and deleting WNK kinases from cells impairs NKCC1/KCC phosphorylation and RVI (Roy et al., 2015b; Zagorska et al., 2007). Thus, WNK kinases play a key role in coordinating the volume recovery response during hypertonic stress.

To undergo activation by hyperosmotic stress, WNK kinases must first overcome a major obstacle – the increase in intracellular chloride and potassium concentrations experienced by cells during volume contraction (Burg, 1995; Cala, 1977; O’Neill, 1999). Increases in intracellular ionic strength, (including [Cl]_i_ and [K+]_i_) potently inhibit WNK kinase activity (Piala et al., 2014; Pleinis et al., 2021). This suggests that an alternate mechanism must be in place that allows the WNKs to sense cell shrinkage, bypass the suppressive effects of ionic strength, and trigger volume recovery via the NKCC1/KCC system.

The mechanisms by which cells detect volume contraction are poorly defined, but likely involve cytoplasmic processes rather than sensors within the plasma membrane (Orlov et al., 2018). Decades ago, investigators proposed that cells adjust their volume not by measuring their cell size *per se*, but rather by monitoring their cytosolic protein concentration (Minton et al., 1992; Parker and Colclasure, 1992). According to this hypothesis, cell shrinkage causes cytoplasmic crowding, which is sensed by signaling pathways that coordinate volume recovery. Crowded intracellular environments may exert such effects by driving the confinement of molecules into compact spaces with altered biochemical activity (Garner and Burg, 1994). For example, macromolecular crowding can induce the formation of biomolecular condensates– functional membraneless microdomains of the cytosol that form via liquid-liquid phase separation (LLPS) (Andre and Spruijt, 2020). Consistent with functional confinement, condensed phases display changes in activity or processing (Shin and Brangwynne, 2017), and thus could coordinate a sensing mechanism that triggers a response to crowding. Indeed, recent work has implicated LLPS as a byproduct of hyperosmotic stress (Cai et al., 2019; Jalihal et al., 2020; Watanabe et al., 2021). Phase separation can occur on rapid timescales; however, it is unclear if biomolecular condensates mediate rapid physiological responses that occur immediately after stress, such as RVI.

Hypertonicity drives WNK1 into cytoplasmic puncta which have long been assumed to be membrane vesicles (Pleiner et al., 2021; Zagorska et al., 2007). Here, we report the unexpected finding that these structures are biomolecular condensates of the WNK signaling pathway that form via LLPS. These liquid WNK1 droplets assemble within seconds, at physiological levels of hypertonic stress, and at native WNK1 protein concentrations. We identify the large WNK1 C-terminal domain as the primary mediator of LLPS, map key regions that confer its phase behavior, and confirm its functional relevance by exploring its phylogenesis and effects on downstream pathway activation, ion transport, and volume regulation. Finally, we show that WNK1 specifically reacts to molecular crowding in vivo, revealing an intrinsic sensing function that allows it to overcome inhibition by chloride and potassium during cell shrinkage.

## Results

### The WNK-SPAK/OSR1 pathway rapidly assembles into dynamic liquid biomolecular condensates following hyperosmotic stress

WNK1 is a ubiquitously expressed serine-threonine kinase that controls cell volume (Roy et al., 2015b). During hyperosmotic stress, WNK1 autoactivates, triggering the downstream phospho-activation of its effector kinases, SPAK (*STK39*) and OSR1 (*OXSR1*), which then directly phosphorylate NKCC1 (*SLC12A2*) and the KCCs (*SLC12A4-7*) (Fig 1A) (Zagorska et al., 2007). While NKCC1 phosphorylation by the WNK-SPAK/OSR1 pathway is activating, phosphorylation of the KCCs by the same pathway is inhibitory (Darman and Forbush, 2002; de Los Heros et al., 2018). Hyperosmotic stress also shifts WNK1 from a diffuse to punctate cytoplasmic distribution (Fig 1B). This change in localization correlates with NKCC1 activation, KCC inhibition, and RVI (Roy et al., 2015b; Zagorska et al., 2007).

**Figure 1.**
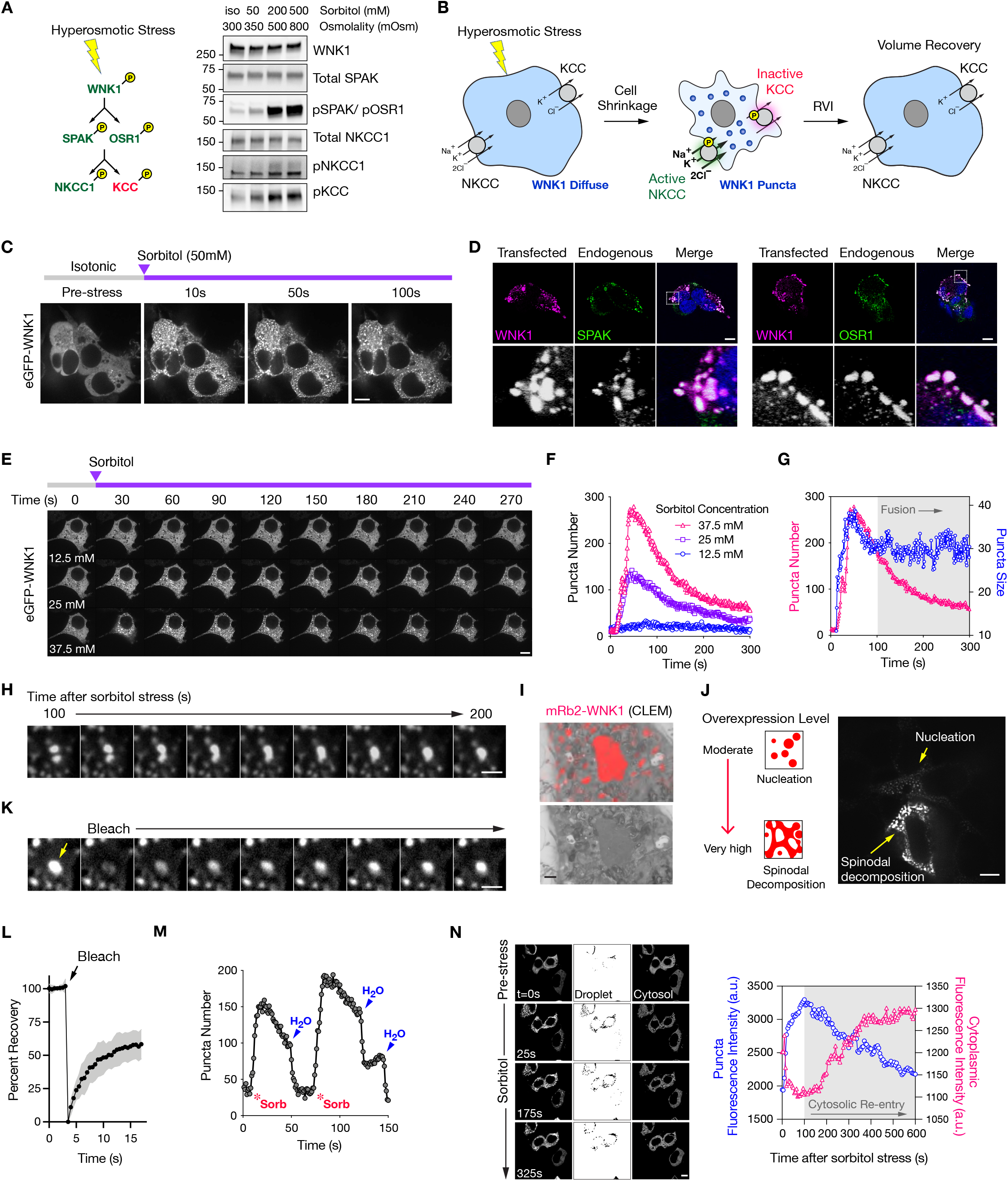
WNK1 forms dynamic liquid-like condensates during hyperosmotic stress. (A) Left: Phosphorylation-dependent regulation of the WNK1-SPAK/OSR1-NKCC/KCC system. Right: Increased endogenous pathway phosphorylation relative to isotonicity (iso), in response to 15min sorbitol stress in HEK cells. See Figure S1A for validation of the total NKCC1 antibody. (B) Hyperosmotic stress is associated with WNK1 puncta formation. NKCC1 and KCC are phosphorylated, activating RVI. (C) Live cell time course of transfected eGFP-WNK1 in HEK cells, subjected to 50mM sorbitol. (D) Fixed IF images of HEK cells transfected with mRuby2-WNK1 subjected to hyperosmotic stress (0.5M sorbitol x 5min), costained for endogenous SPAK and OSR1. (E-F) Representative live cell time course of a HEK cell expressing eGFP-WNK1, subjected to a low-level sorbitol stress series (12.5mM-37.5mM), with corresponding quantification of puncta number over time. (G) Comparison of puncta size and number in eGFP-WNK1 expressing cells subjected to 37.5 mM sorbitol, as shown in (E). 100s into the time course, puncta size stabilized while total number falls, consistent with fusion. (H) Fusion between two eGFP-WNK1 puncta 100s after 37.5mM sorbitol treatment. Bar= 2µm. (I) CLEM of a WNK1 condensate in fixed mRuby2-WNK1 transfected cells subjected to 0.1M sorbitol x 15min. Bar= 1µm. (J) Live cell imaging still of two cells expressing moderate and very high levels of eGFP-WNK1, subjected to 50mM KCl, demonstrating nucleation and spinodal decomposition, respectively. Still image from Video S2. (K-L) FRAP recovery images and corresponding recovery curve of eGFP-WNK1 condensates in cells subjected to 50mM sorbitol. Bar= 2µm. (M) Reversibility of WNK1 puncta number over time in eGFP-WNK1 transfected cells subjected to osmotic challenges with 50 mM sorbitol, followed by water. Bar= 2µm. (N) Quantification of eGFP droplet and cytosolic fluorescence intensity over time in transfected cells. All bars = 10µm except where indicated.

The role of these puncta in WNK signaling has been unclear; thus, we sought to understand their dynamics and functional relevance to the volume recovery response. In live cell imaging studies, transiently expressed mammalian (rat) WNK1 shifted to cytoplasmic puncta with modest hyperosmotic stress (50mM sorbitol; 350mOsm) (Fig 1C), and the foci colocalized with SPAK and OSR1 (Fig 1D), suggesting they physiologically regulate WNK signaling. The puncta were visible with sorbitol stresses as low as 25mM (325 mOsm), and the number of puncta that formed per cell increased with higher degrees of osmotic stress (Fig 1E & 1F). Other hyperosmotic stressors exerted similar effects on WNK1 puncta formation (Fig S1B & S1C). In contrast, low Cl- hypotonic stress — a maneuver commonly used to evaluate WNK kinase activity — had no apparent effect on WNK distribution (Fig S1D).

The WNK1 puncta appeared immediately after stress and grew steadily in size, but after 100s, their size stabilized despite a steady decrease in number (Fig 1G). This was due to the fusion of individual puncta (Fig 1H, Video S1). As the puncta merged, they behaved like liquids, resolved into round structures (Fig 1H and Video S1), wetted against intracellular surfaces (Fig S1E, Video S1), and were membraneless and electron hyperdense in correlative light and transmission electron microscopy (CLEM) experiments (Fig 1I). Collectively, these features are typical of condensates that form via LLPS. At modest WNK1 expression levels, the puncta underwent nucleation and growth (Fig 1F, 1G, & 1J). In cells expressing WNK1 at very high levels, hyperosmotic stress triggered its instantaneous demixing into networks that appeared bicontinuous, consistent with spinodal decomposition (Fig 1J & Video S2) (Shimobayashi et al., 2021). Concentration-dependent nucleation and spinodal decomposition are hallmarks of phase-separating systems (Alberti et al., 2019)

Consistent with dynamic liquid phase behavior, the WNK1 droplets exhibited rapid recovery kinetics in photobleaching experiments (Fig 1K & 1L and Video S1) and dissolved within seconds when the hypertonic stress was quenched with water (Fig 1M). In addition, changes in WNK1 fluorescence within and outside of the condensed phase were linked, as the reduction in droplet fluorescence during later stages of the post-stress time course was associated with a reciprocal increase in diffuse phase signal (Fig 1N). Thus, condensed material eventually leaves the droplet phase and re-enters the cytosol.

### Endogenous WNK1 forms hyperosmotic stress-induced condensates

To determine if these findings are relevant at native levels of WNK1 expression, several approaches were employed. First, we evaluated endogenous WNK1 localization in fixed HEK- 293 cells subjected to hyperosmotic stress, using a validated WNK1 antibody (Fig S2A & B). Osmotically stressed cells exhibited high-intensity WNK1 puncta that nucleated above optical diffraction limits (Fig 2A & B) and became larger with increasing degrees of hypertonicity (Fig 2C & D). Similar findings were observed in U2OS and GBM43 glioma cells (Fig S2C). In all three cell lines tested, endogenous WNK1 was occasionally present in small high intensity puncta under iso-osmotic conditions (Fig 2C, 2D, S2C). This suggests that low-level WNK1 puncta formation could contribute to its endogenous baseline activity during isotonicity.

**Figure 2.**
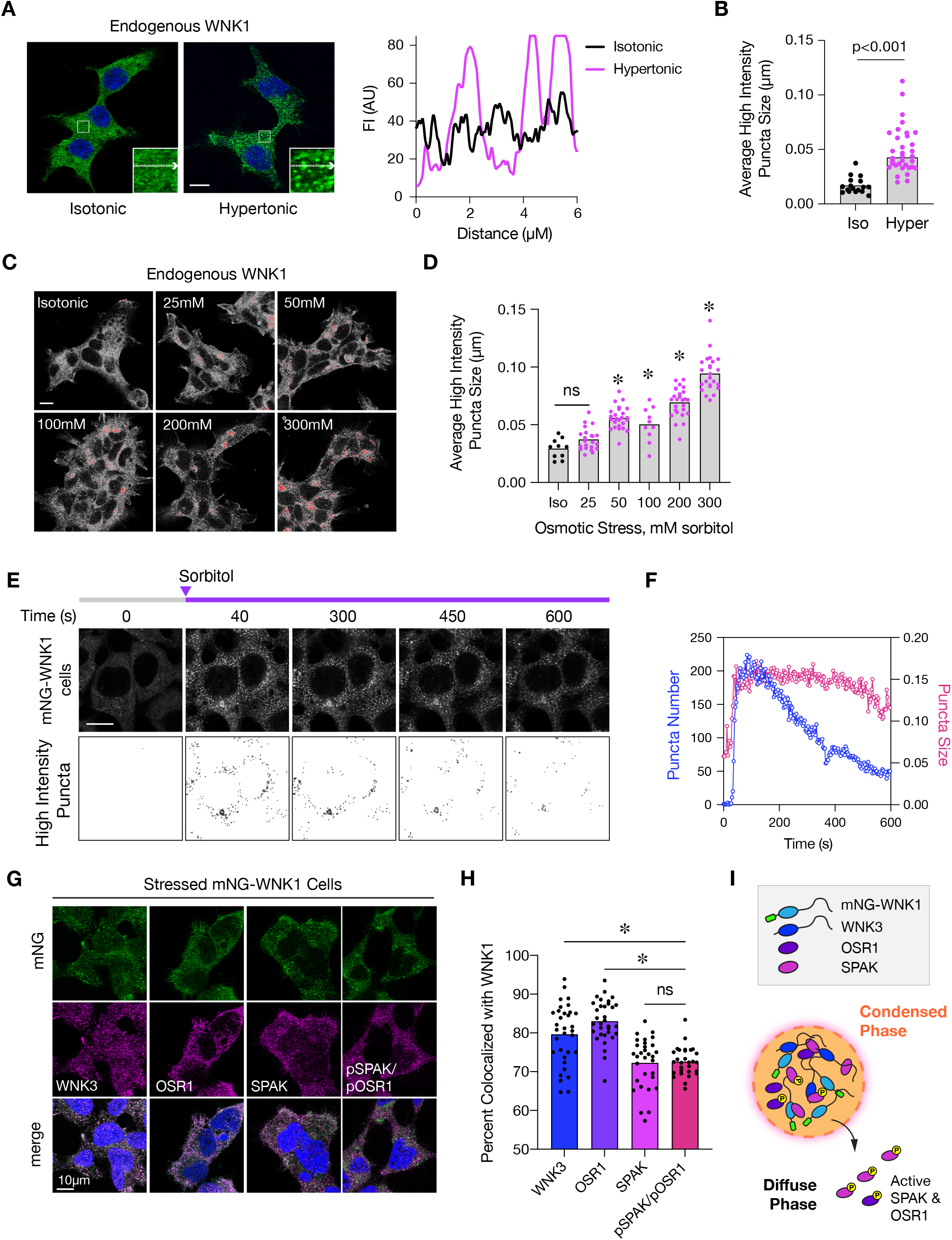
Endogenous WNK1 forms hyperosmotic stress-induced condensates. (A) Fixed WNK1 IF images in HEK cells under isotonic and hypertonic conditions (300mM sorbitol x 5min). Line scans on the right depict fluorescence intensities (FI) along the inset arrows on the left. (B) Quantification of average high intensity puncta size/confocal field in HEK cells under iso- (n=17 fields) and hypertonic conditions (n=34 fields). p<0.0001 by unpaired T-test. High intensity signal was defined by manual thresholding in FIJI, which was kept constant across all conditions analyzed. (C) Fixed IF images of endogenous WNK1 in HEK cells subjected to various degrees of hypertonic stress (mM sorbitol). Thresholded high intensity signal is shown in red. (D) Quantification of average high-intensity puncta size/ confocal field in (C). n=10-26 confocal fields per condition; *p<0.0008 by one-way ANOVA, Dunnett’s post-hoc test vs isotonic control. (E-F) Representative live cell time course of mNG signal in mNG-WNK1 knock-in cells subjected to 50mM sorbitol, with corresponding quantification of high-intensity puncta number and size over time. (G-H) Colocalization of WNK-SPAK/OSR1 pathway components with mNG-WNK1 in fixed cells subjected to 300mM sorbitol. N=29-33 confocal fields imaged per immunostaining condition. *p<0.0001 by one-way ANOVA, Dunnett’s post-hoc test vs pSPAK/pOSR1. (I) While WNK3 is strongly colocalized with WNK1 in condensates, active SPAK/OSR1 are more localized in the cytosol. All bars = 10µm.

To assess native WNK1 condensate dynamics, we employed CRISPR/Cas9 technology to introduce an in-frame N-terminal mNeonGreen fluorescence tag into the first exon of WNK1 (Fig. S2D). This modification had no effects on WNK1 expression, or on its ability to activate SPAK/OSR1 (Fig. S2E). Though the mNG-WNK1 puncta were smaller than those observed during overexpressed conditions, they nucleated seconds after hyperosmotic stress and underwent fusion (Video S3). Similar to the overexpressed WNK1 condensates, the fusion events resulted in high-intensity mNG-WNK1 puncta that steadily decreased in number but maintained a relatively stable size over a time course of minutes (Fig 2E, F). Consistent with a physiologic role in WNK signaling, mNG-WNK1 puncta contained the osmotic stress-responsive paralog WNK3 (Akella et al., 2021; Pacheco-Alvarez et al., 2020), SPAK, and OSR1 (Fig 2G). The WNK3 and OSR1 signals were 80% colocalized with mNG-WNK, while the active phosphorylated forms of SPAK and OSR1 (pSPAK/pOSR1) were less colocalized at 70%. This suggests that active SPAK and OSR1 can leave the condensed phase following activation (Fig 2H & I).

To further characterize the endogenous mNG-WNK1 puncta, we performed high resolution colocalization studies between mNG-WNK1 puncta and subcellular markers using confocal microscopy with adaptive deconvolution. The puncta did not colocalize with endosomes (EEA1) or lysosomes (LAMP2) (Fig S3A & B). In contrast to an earlier report (Zagorska et al., 2007), we also did not observe significant colocalization between mNG-WNK1 and the AP-1 adaptor complex. However, consistent with prior work (Pleiner et al., 2021; Zagorska et al., 2007), the stress-induced WNK1 puncta colocalized strongly with clathrin heavy chain (Fig S3A & B). Hypertonic stress suppresses endocytosis, coated pit formation, and releases clathrin heavy chain from membranes into the cytosol (Hansen et al., 1993; Heuser and Anderson, 1989). Since clathrin heavy chain also physically associates with WNK1 (Pleiner et al., 2021), these findings indicate that WNK1 sequesters clathrin within membraneless condensates during cell shrinkage. Overexpressed WNK1 also colocalized strongly with clathrin heavy chain during hyperosmotic stress, indicating that the condensates characterized in Figure 1 and natively expressed mNG- WNK1 puncta share the same features (Fig S3C). Further supporting the membraneless nature of mNG-WNK1 puncta, they partially colocalized with EMC2, a soluble subunit of the ER membrane protein complex that binds to WNK1 in the cytosol (Pleiner et al., 2021). Notably, the WNK1 condensates failed to colocalize with TIAR-1, eIF4e, and DCP2 (Fig S3A & B), indicating that they are distinct from stress granules and P-bodies.

### The intrinsically disordered C-terminal domain mediates WNK1 phase separation

Homo- and heterotypic molecular interactions drive protein phase behavior (Riback et al., 2020). For example, transient multivalent interactions between SH3 domains and binding partners harboring proline-rich motifs can trigger LLPS (Li et al., 2012). Alternatively, LLPS often requires weak associations between intrinsically disordered regions (IDRs) of low sequence complexity that contain prion-like domains (PLDs) (Shin and Brangwynne, 2017). WNK1 harbors numerous proline-rich motifs that bind to SH3 and WW domain partners (He et al., 2007; Roy et al., 2015a), and disorder prediction analysis revealed that nearly the entire WNK1 polypeptide sequence outside of the structured kinase domain is disordered and of low complexity (Fig 3A). This includes the large C-terminal domain (CTD), which harbors two coiled coil domains adjacent to low-complexity PLDs. Thus, WNK1 contains multiple features that may confer its phase behavior.

**Figure 3.**
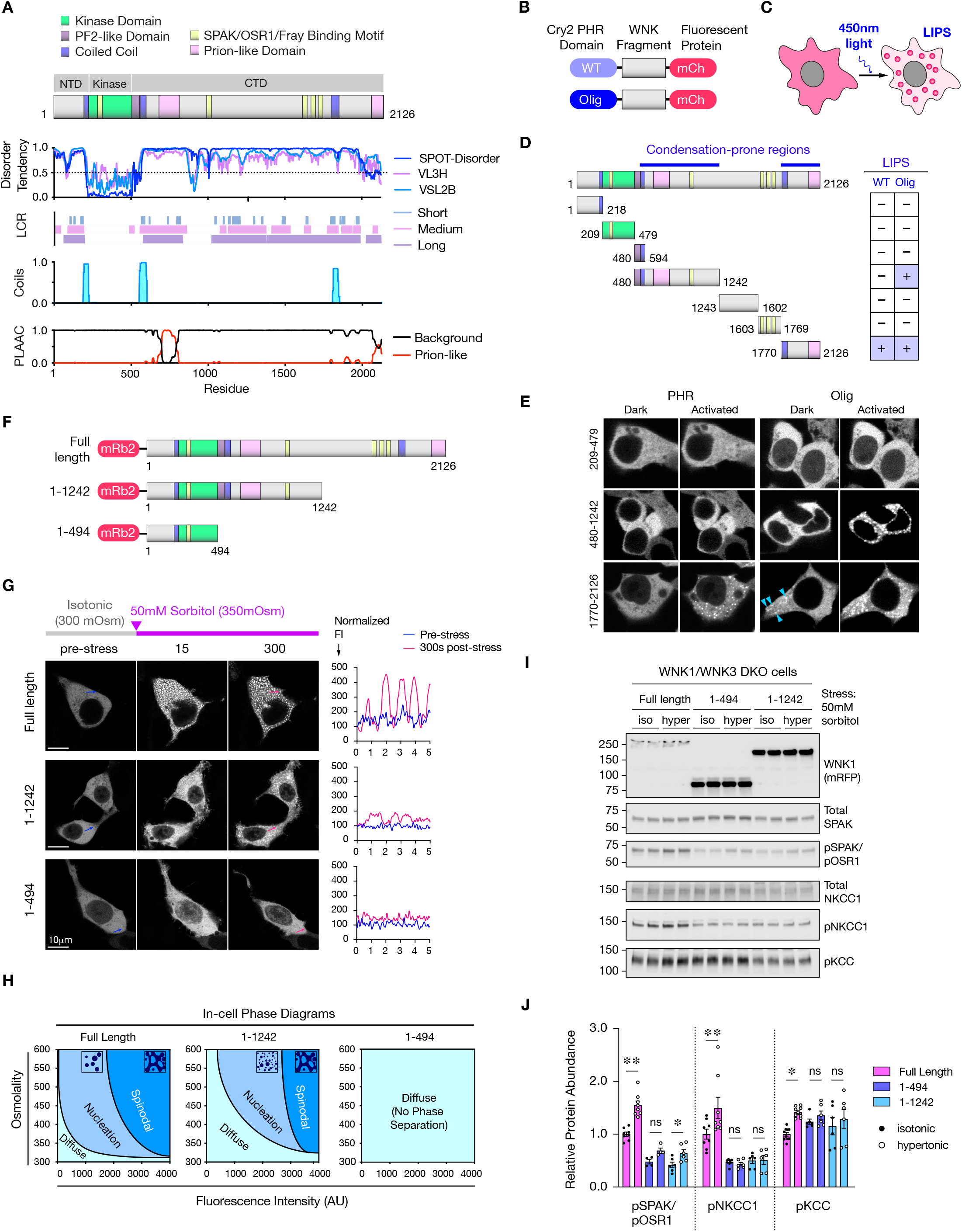
WNK1 phase separation is mediated by its intrinsically disordered CTD. (A) Mammalian (Rat) WNK1 domain structure, with key domains color-coded on top. The kinase domain is flanked by an N-terminal domain (NTD) and a large C-terminal domain (CTD). Below is a to-scale bioinformatic analysis for protein disorder, low complexity regions (LCR) of short, medium, and long trigger window length, Coiled-coil domains, and Prion-like regions. (B-C) Schematic of optogenetic WNK1 constructs, consisting of WT or E490G mutant (“Olig”) PHR domains, fused in-frame to mCherry-tagged WNK1 fragments. Blue light activation of LLPS- prone fragments triggers light-inducible phase separation (LIPS). (D) Analysis of PHR-WNK1-mCh fusions. Condensation-prone fragments are depicted with at “+”. Two fragments C-terminal to the kinase domain underwent LIPS. (E) Still image examples from live cell LIPS experiments. Two fragments underwent LIPS (480- 1242, Olig only; 1770-2126; WT and Olig). 1770-2126 also exhibited spontaneous dark state aggregation when fused to Olig (arrowheads). In contrast, 209-479 (the kinase domain) did not undergo LIPS. (F) Full length and truncated N-terminal mRuby2-tagged WNK1 constructs, (G) Representative live cell images of cells expressing the constructs in (F), pre- and post- osmotic stress. Line scans on the right depict normalized pre- and post-stress fluorescence intensities (FI) along the arrows depicted on the right. (H) In-cell phase diagrams depicting the phase behavior of the constructs in (F), as a function of fluorescence intensity (concentration) and extracellular osmolality. (I-J) SPAK/OSR1/NKCC1/KCC activation in WNK1/WNK3 DKO cells expressing the constructs in (F), under isotonic and hypertonic conditions (50mM sorbitol x 30min). p< *0.05 or **0.01 by one-way ANOVA, Sidak’s multiple comparisons test.

To screen for regions of WNK1 that are prone to phase separation, we employed an optogenetic approach that takes advantage of the photolyase homology region (PHR) of *Arabidopsis thaliana* Cry2, which transiently self-associates upon exposure to blue light. Fusing a condensation-prone sequence to the Cry2-PHR domain triggers light-inducible phase separation (LIPS) (Figs 3B & 3C) (Mann et al., 2019; Shin et al., 2017). Cry2Olig, an engineered PHR variant, exhibits higher sensitivity to light-induced clustering (Taslimi et al., 2014) (Fig 3B). We screened for regions in WNK1 that undergo strong or weak LIPS by generating WNK1 fragments fused to either the wild-type (WT) PHR domain of Cry2, or to Cry2Olig (Fig 3D). This approach mapped two areas within the WNK1 CTD that underwent LIPS. One fragment encoding the extreme C-terminal end (1770- 2126) was strongly prone to phase separation, as both the WT and Cry2Olig fusions underwent LIPS (Fig 3D & 3E, Video S4). In addition, rare spontaneous puncta were appreciable with the Cry2Olig construct in the dark state, further supporting its highly condensation-prone nature (Fig 3E, arrowheads). A second more proximal region (480-1242) condensed less strongly, as only the Cry2Olig fusion underwent LIPS (Fig 3D & 3E). Notably, both condensation-prone fragments are low complexity IDRs within the WNK1 CTD that contain coiled coil domains and prion-like signatures. Other regions of WNK1, such as the kinase domain (209-479) failed to undergo LIPS in either the WT PHR or Cry2Olig constructs (Fig 3D & 3E, Video S4).

To evaluate whether the WNK1 CTD mediates osmotic stress-induced LLPS, we compared the phase behavior of full length WNK1 to two fragments: (1-1242), containing the proximal condensation-prone region, and (1-494), which lacks the entire CTD (Fig 3F). Given the strong dependence of phase behavior on protein concentration (Alberti et al., 2019), cells with similar levels of pre-stress fluorescence were compared. While the 1-1242 construct underwent LLPS, it failed to efficiently clear from the cytosol, resulting in smaller less well-defined condensates compared to the full-length protein (Fig 3G, Video S5). The 1-494 construct did not phase separate, instead maintaining a diffuse cellular signal following osmotic stress (Fig 3G, Video S5).

To get a better sense of how the phase behavior of these constructs is tied to WNK1 expression levels, we generated partial in-cell phase diagrams that estimate in vivo LLPS at different degrees of hyperosmotic stress as a function of pre-stress fluorescence intensity, a surrogate for protein concentration (Fig S4A & B). Both the full-length and 1-1242 constructs formed stress-induced nucleated and spinodal assemblies (Fig S4A). However, compared to full length WNK1, the 1- 1242 construct required higher levels of protein expression and hyperosmotic stress to uniformly cross the transition threshold from a diffuse to nucleated state (a phase boundary referred to as the binodal (Shimobayashi et al., 2021); Fig S4B & 3H). On the other hand, the spinodal phase boundary (i.e., the threshold at which a protein undergoes spinodal decomposition instead of nucleation) was roughly similar for the two constructs, particularly at hyperosmotic stresses below 400mOsm. Thus, compared to full-length WNK1, the binodal and spinodal curves defining the phase behavior of the 1-1242 construct were contracted, resulting in a narrowing of the nucleation window within the physiologic range of hyperosmotic stress (Fig 3H). In contrast to these two constructs, the 1-494 fragment did not form visible condensates at any of the osmotic stresses tested, and thus appeared to exist in a one-phase regime.

If efficient WNK nucleation into droplets augments kinase activation, the LLPS-deficient 1-494 and 1-1242 constructs should exhibit less activation than the full-length protein. To test this, we performed biochemical assessments of the WNK-SPAK/OSR1 pathway at physiologic levels of hyperosmotic stress (350mOsm). To eliminate the influence of endogenous osmotically- responsive WNK kinases, we conducted these studies in gene-edited WNK1/WNK3 double knockout (DKO) cells (Fig S5). In DKO cells expressing transfected full-length WNK1, 50mM sorbitol triggered immunodetectable phosphoactivation of SPAK/OSR1 and downstream phosphorylation of NKCC1 and KCCs (Fig 3I & 3J). In contrast, there was little to no increase in phosphorylation of SPAK/OSR1, NKCC1, or KCC in DKO cells transfected with the 1-1242 or 1-494 constructs (Fig 3J). Combining these biochemical studies with the live cell imaging data, these results collectively indicate that the entire C-terminal domain is necessary for WNK1 to drive optimal SPAK/OSR1 activation and downstream NKCC1/KCC phosphorylation via phase separation.

### C-terminal coiled-coil domains augment WNK1 phase behavior and activity

Coiled coil domains (CCDs) are often present within proteins that undergo LLPS (Ford and Fioriti, 2020). Because CCDs reside within both regions within the WNK1 C-terminal domain that confer its phase behavior (Figs 3A), we sought to determine their importance. CCs are formed by amino acid heptad repeats (denoted by residues *a* through *g*). Positions *a* and *d* define the CC core and are frequently occupied by hydrophobic residues, or by “ambivalent hydrophobes” such as glutamine (Fig 4A) (Fiumara et al., 2010; Sodek et al., 1972). Hydrophobic and glutamine residues were present in the *a* and *d* registers of both C-terminal WNK1 CCDs (Fig 4B). In the C-terminal CC (CT-CC), the *a*-*d* frame also contained a known “HQ” signature which facilitates WNK-WNK interactions (Thastrup et al., 2012). We substituted prolines into these predicted key residues to disrupt CC structure without influencing regional disorder tendency (Fig 4B & 4C). This allowed us to specifically evaluate the CCDs independently of IDR function.

**Figure 4.**
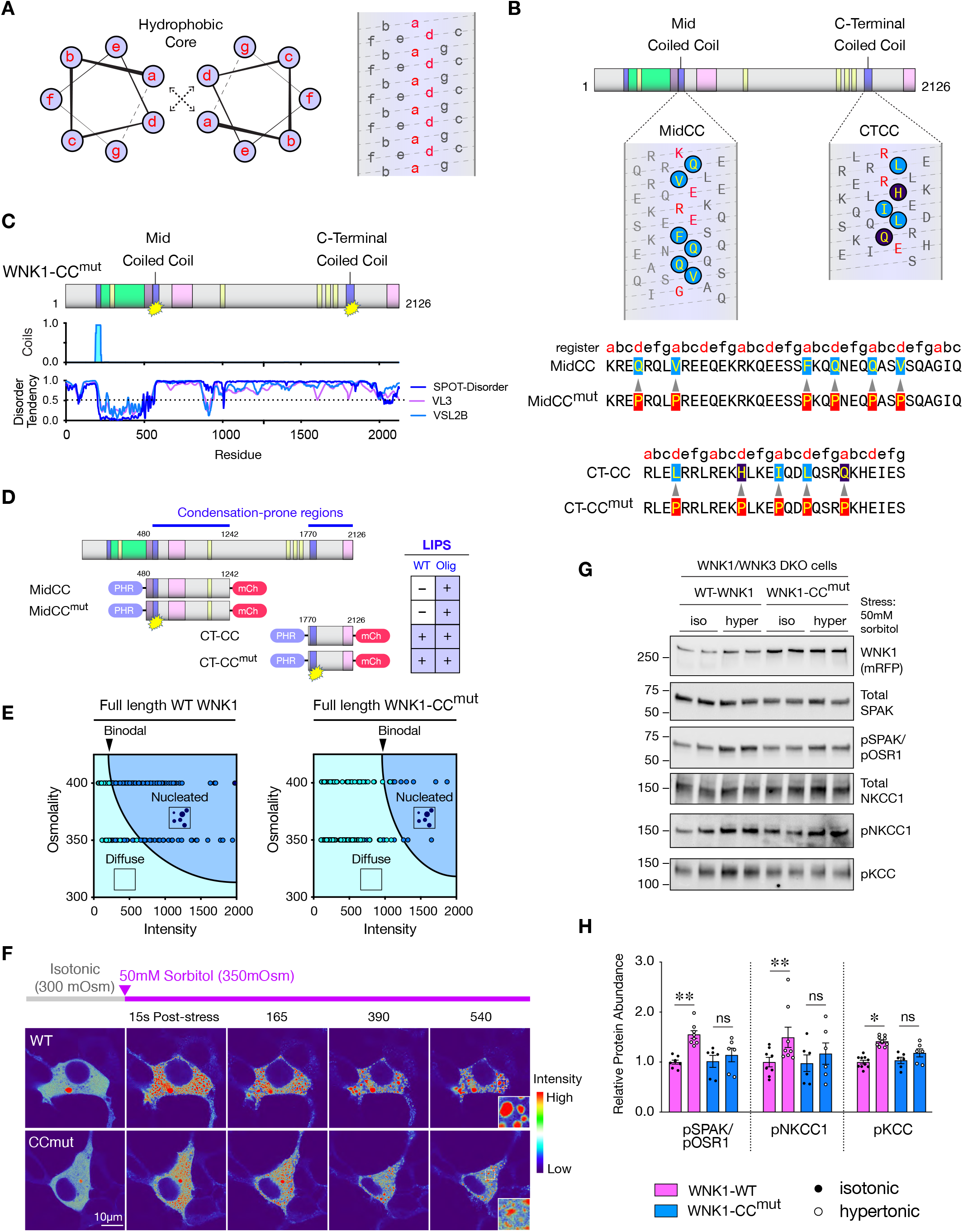
C-terminal coiled coil domains augment WNK1 phase behavior and activity. (A) Helical wheel and net diagrams of coiled coil (CC) heptad repeats. (B) Helical net diagram of the MidCC and CTCC domains in rat WNK1. Hydrophobic and Q residues within the *a/d* frame are circled. An “HQ” signature important for CTCC interactions resides within the *a/d* frame. To disrupt CC function, key proline substitutions were introduced into the *a/d* register, as indicated. (C) WNK1 double coiled-coil mutant (WNK1-CC^mut^). The proline substitutions disrupted predicted CC structure without affecting disorder tendency. (D) Proline mutagenesis of the midCC and CTCC did not affect LIPS in the Cry2-PHR system. (E) Mutation of the midCC and CTCC in the context of full-length WNK1 (WNK1-CC^mut^) introduced a rightward shift of the binodal at low levels of hypertonic stress. (F) LLPS in sorbitol-stressed HEK cells expressing mRb2-tagged WT Full-length WNK1 and WNK1-CCmut. Despite similar pre-stress expression levels, the CC mutant formed smaller short- lived condensates. Differential intensities were pseudocolored using the Thermal lookup table in FIJI. (G-H) Pathway activation in WNK1/WNK3 DKO cells expressing WT-WNK1 or WNK1-CC^mut^, under isotonic and hypertonic conditions (50mM sorbitol x 30min). p< *0.05 or **0.01 by one-way ANOVA, Sidak’s multiple comparisons test.

Optogenetic assessment of phase separating WNK1 fragments containing the CCDs revealed no effect of the proline mutations on LIPS (Fig 4D). However, in comparative in-cell phase diagrams, a full-length double mutant construct containing proline substitutions in both the MidCC and CT- CC (WNK1-CC^mut^; Fig 4C) exhibited a rightward shift of the binodal phase boundary (Fig 4E). Furthermore, in WNK1-CC^mut^ expressing cells that formed WNK condensates, disruption of the C-terminal CCDs resulted in smaller puncta that were more short-lived compared to WT-WNK1. Consistent with reduced WNK1 retention within condensates, this was associated with higher fluorescence intensity in the diffuse phase (Fig 4F). Thus, mutation of the CCDs impaired WNK1’s ability to efficiently form large droplets that increase in size and decrease in number as they fuse and extract material from the diffuse phase, a process referred to as coarsening (Hyman et al., 2014). In comparative immunoblots with full length WT-WNK1, coiled-coil domain disruption impaired downstream SPAK/OSR1, NKCC1, and KCC phosphorylation (Fig 4G & 4H). Collectively, these findings indicate that though the C-terminal CCDs are not necessary for LLPS, they facilitate optimal WNK1 nucleation and pathway activation within condensates.

### Phase behavior of the WNK1 CTD is evolutionarily conserved despite poor sequence identity

We wondered if the disorder features present in rat WNK1 are present in other WNK kinases. Thus, we analyzed four WNKs with poorly alignable sequences outside of the kinase domain that mediate similar functions with regards to osmotic stress responsiveness and/or ion transport: human WNK1 (hWNK1), human WNK3 (hWNK3), *Drosophila melanogaster* WNK (dmWNK), and *Caenorhabditis elegans* WNK (ceWNK). Despite their poor sequence identity, the disorder tendency plots and prion-like signatures for these WNKs are nearly identical (Fig S6A). To further explore sequence relationships in WNK IDRs that likely exhibit phase behavior, we analyzed the amino acid composition of WNK CTDs across evolution, extending from humans to unicellular eukaryotes (Fig S6B, Table S1). Mammalian WNK kinases are enriched in prolines and serines, which comprise approximately 25% of the amino acids in the disordered CTD. This compositional bias extends to freshwater vertebrate fish. However, in many organisms extending from invertebrates down to protists, glutamine is the preferred amino acid. Glutamine and serine regulate the material properties of condensates in a similar manner (Wang et al., 2018). Thus, the WNK CTD underwent a permissive Q-to-P/S switch in compositional bias during early vertebrate evolution that conserved its regional disorder tendency, likely to preserve its phase behavior. This suggests that the ability of WNK kinases to form condensates is an ancient, fundamental property.

To test this further, we turned to the *Drosophila melanogaster* WNK kinase. dmWNK is a central regulator of ion transport in the Malpighian tubule (MT), an ancient but conserved model of the mammalian nephron (Rodan, 2018). Within this system, dmWNK triggers NKCC phosphoactivation via the SPAK homolog Fray. While the rat WNK1 CTD bears a P/S-rich composition present in other mammalian WNKs, the dmWNK CTD is Q-enriched, as is the case for many invertebrates (Fig S6B). These glutamines cluster in poly-Q tracts that reside in the proximal and distal regions of the C-terminal domain – PLDs that correspond to regions in rat WNK1 that were identified to be critical for LLPS (Fig 5A). Though poly-Q prion-like sequences are classically considered to be mediators of pathological protein aggregation, they can facilitate physiological stress-induced phase transitions (Franzmann et al., 2018).

**Figure 5.**
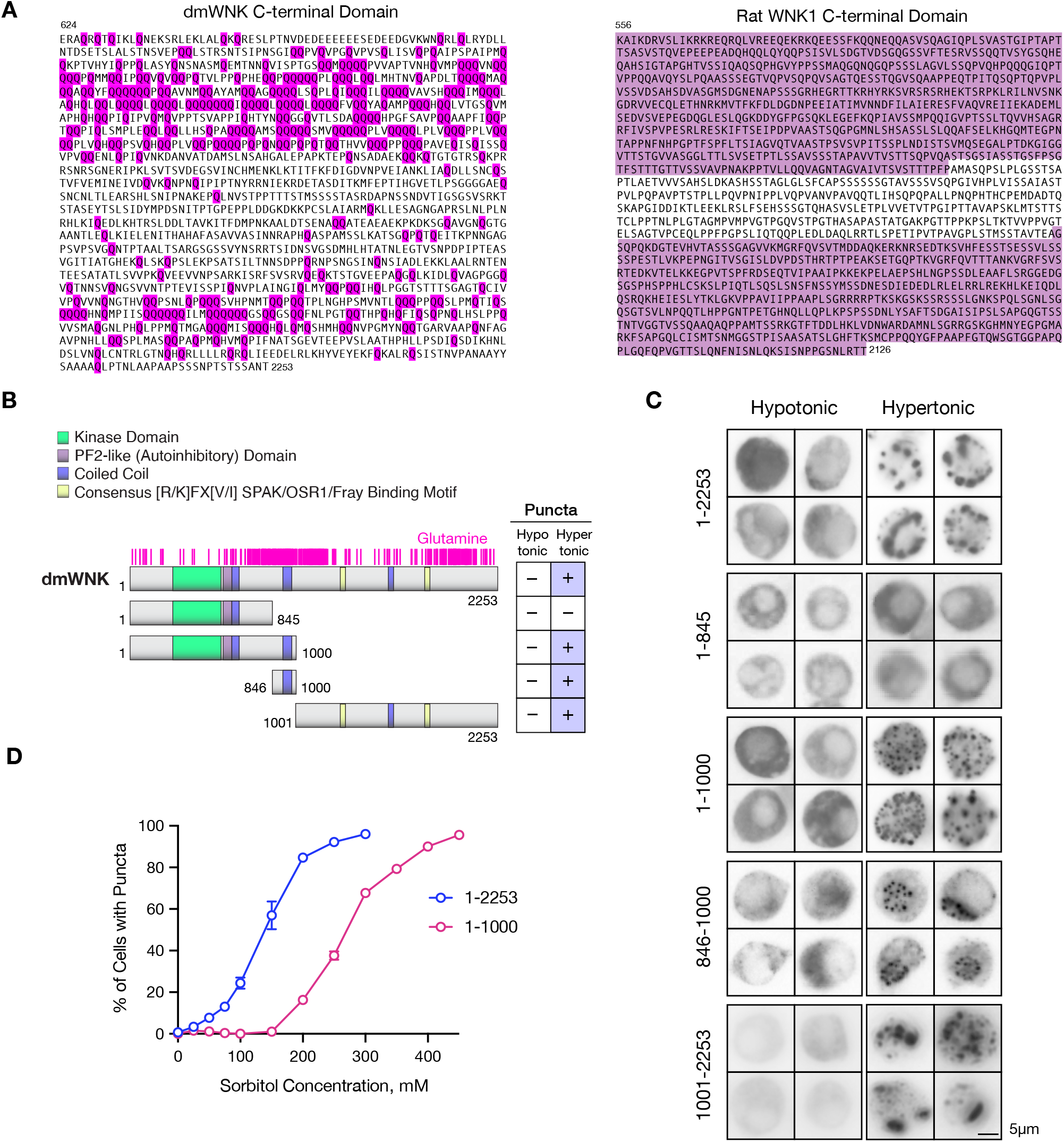
Evolutionary conservation of WNK phase behavior in *Drosophila*. (A) Amino acid sequences of the rat WNK1 and dmWNK CTDs. Glutamines in dmWNK are shown in magenta. Condensation-prone regions in rat WNK1 identified in Figure 3 are highlighted in lilac. (B-C) dmWNK deletion analysis. Diagram of eGFP-tagged dmWNK truncation constructs, with glutamine residues shown in magenta. Apart from dmWNK 1-845, all constructs tested formed puncta during hypertonic stress. (D) Compared to full-length dmWNK, a construct containing only the N-terminal part of the CTD triggered puncta formation in fewer cells at a given level of hyperosmotic stress.

Given these observations, we asked whether dmWNK phase separates during hyperosmotic stress. In live cell imaging studies in *Drosophila* S2 cells overexpressing GFP-tagged dmWNK, the full-length construct (1-2253) behaved similarly to mammalian WNK1, shifting to condensates during hyperosmotic stress (Video S6, Fig 5C). A deletion analysis confirmed that like rat WNK1, dmWNK harbors two zones within its CTD that are primarily responsible for its phase behavior. One of these (846-1000) is a Q-rich region in the middle of the protein that harbors a CCD, while the other is contained within the more distal region of the CTD (1001-2253) (Fig 5B & C). While a dmWNK fragment lacking this distal region (1-1000) still formed condensates (Fig 5C), it exhibited LLPS in fewer cells compared to the full-length protein at a given level of osmotic stress (Fig 5D). Thus, like mammalian WNK1, the entire intrinsically disordered CTD is required for optimal LLPS. These findings reinforce evidence that WNK phase behavior is highly conserved and encoded within specific regions of the CTD.

### The WNK1 CTD is required for NKCC1 transport and RVI

To determine whether the WNK1 CTD influences RVI, we performed cell volume measurements using high-speed live cell resonant scanning confocal microscopy to capture rapid volumetric changes within the phase separation time frame. RVI is classically mediated by two transport processes: a cariporide-sensitive component involving coupled sodium/proton and chloride/bicarbonate exchange and a bumetanide-sensitive component mediated by the NKCC1/KCC system (Fig 6A) (Hoffmann et al., 2009). To rule out compensation by the sodium/proton exchanger NHE1, all volume measurements were conducted in the presence of cariporide. Following exposure to 50mM sorbitol, cells expressing full-length rat WNK1 on a WNK1/WNK3 DKO background exhibited a rapid reduction in cell volume, followed by a modest linear recovery over time, consistent with classical RVI (Fig 6B & 6C). In contrast, cells expressing the CTD-deficient rat 1-494 construct exhibited a sluggish RVI response following acute cell shrinkage, with a rundown of cell volume that continued for 5 minutes post-stress prior to showing evidence of recovery (Fig 6B & C). In cells expressing full-length WNK1, preincubation with the NKCC1 inhibitor bumetanide blunted the RVI response within the first five minutes of the recovery time course (Fig 6D). In contrast, cells expressing WNK1 1-494 exhibited similar RVI curves in the presence and absence of bumetanide (Fig 6E). Thus, DKO cells expressing the full length WNK1 construct mounted a bumetanide-sensitive NKCC1-mediated RVI response that was most evident during the initial recovery period, while cells expressing the LLPS-deficient 1-494 construct did not (Fig 6F).

**Figure 6.**
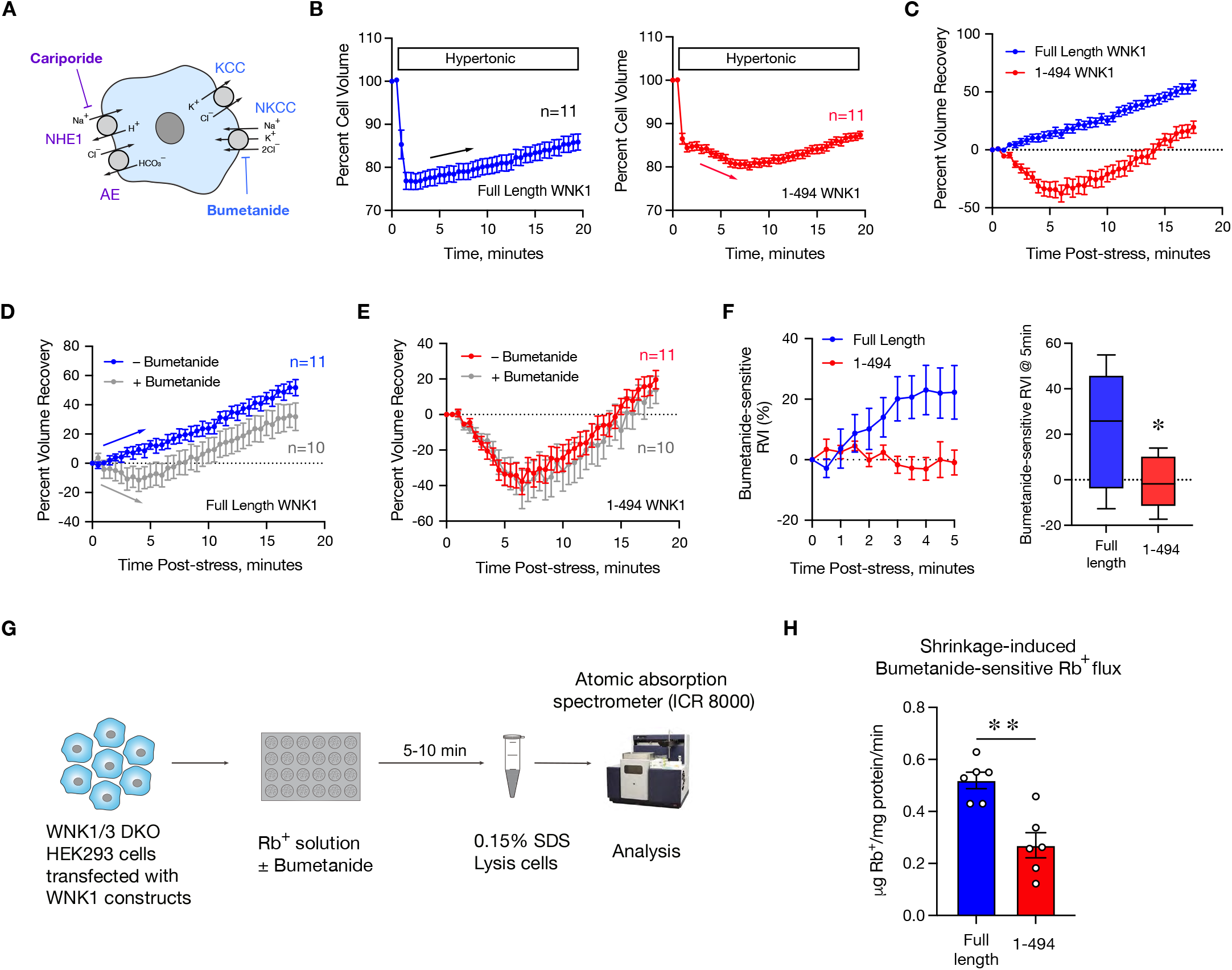
WNK1 CTD augments NKCC1-dependent RVI. (A) Cariporide-sensitive NHE1/AE and bumetanide-sensitive NKCC1/KCC RVI systems. RVI measurements were conducted in the presence of cariporide to rule out NHE1-dependent effects. (B) Cell volume changes in WNK1/WNK3 DKO cells, transfected with full-length WNK1 or 1-494 WNK1. (C) RVI curves derived from the cell volume measurements in (B), expressed as % volume recovery following cell shrinkage. (D-E) RVI curves in DKO cells transfected with full-length WNK1 or 1-494 WNK1, +/- Bumetanide. (F) Bumetanide-sensitive RVI, calculated as the difference between the +/- bumetanide curves in (D) & (E). *p=0.0320 for Full-length WNK1 vs 1-494 WNK1 expressing DKO cells @ 5min post stress, unpaired T-test. (G) Schematic diagram of Rb flux experiment. (H) Shrinkage-induced Bumetanide-sensitive Rb flux in WNK1/WNK3 DKO cells transfected with full-length WNK1 or 1-494 WNK1. N=6 sets of flux measurements per condition. **p=0.0015 by unpaired T-test.

To determine the functional importance of the disordered WNK1 CTD on NKCC1 transport, we used atomic absorption spectroscopy to measure bumetanide-sensitive influx of the K+ congener rubidium (Rb+) under isotonic and hypertonic conditions in WNK1/WNK3 DKO cells expressing either full length or 1-494 WNK1 (Fig 6G). A shift in osmolality from near isotonicity (310 mOsm) to hypertonicity (370 mOsm) was associated with an increase in bumetanide-sensitive Rb+ influx. However, this increase was blunted in cells expressing 1-494 WNK1 compared to full-length WNK1 (Fig 6H).

### WNK1 is a molecular crowding sensor

Macromolecular crowding commonly triggers protein phase separation and has also been proposed to be a stimulus for RVI (Andre and Spruijt, 2020; Colclasure and Parker, 1991). Thus, we hypothesized that the formation of bioactive WNK1 condensates may be the manifestation of a crowding-sensing mechanism that triggers the RVI response. To test this, we designed an experiment that augments cytoplasmic molecular crowding without causing cell shrinkage. Ficoll is a molecular crowding agent that is commonly used to trigger protein phase separation in vitro (Andre and Spruijt, 2020). We reasoned that cytoplasmic Ficoll microinjection should increase the degree of cytosolic crowding while simultaneously increasing cell size. If a protein phase separates under such conditions, it could be reasoned that it does so because of molecular crowding rather than shrinkage (Fig 7A).

**Figure 7.**
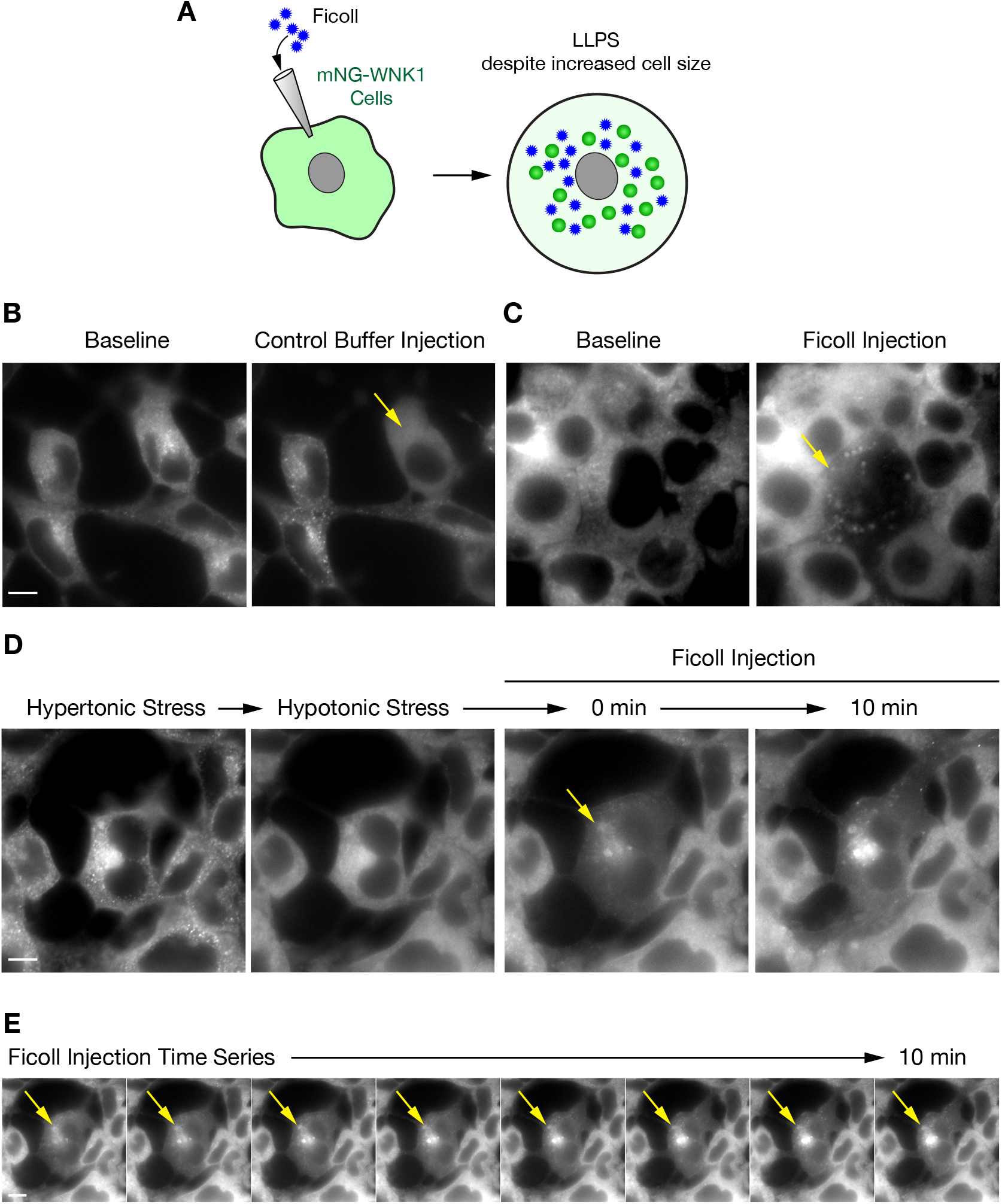
WNK1 is a molecular crowding sensor. (A) Ficoll microinjection experiment. (B) Effect of KCl control buffer injection into mNG WNK1 cells (C) Microinjection of Ficoll-containing buffer increased mNG-WNK1 puncta formation. (D) Effect of Ficoll microinjection following hypotonic cell swelling. (E) 10min timecourse following Ficoll injection. Crowding agent induced localized long-lived puncta formation All bars = 10µm.

For these studies, we used mNG-WNK1 cells, which clonally express fluorescently tagged WNK1 at native levels. As noted in Fig 2E, these cells exhibit low-grade WNK1 condensation under isotonic conditions, resulting in a granular appearing cytoplasm with partial localization of mNG fluorescence in small puncta (Fig 7B). Following the microinjection of a control vehicle buffer lacking crowding agent, cells expanded and the mNG signal became diffuse and homogeneous, consistent with condensate dissolution due to cytoplasmic dilution (Fig 7B). In contrast, Ficoll microinjection triggered the formation of large WNK1 condensates which extracted WNK1 from the surrounding dilute phase (Fig 7C). Introduction of Ficoll into swollen cells that were precleared of puncta by hypotonic stress was associated with the localized collection of long-lived condensates that coarsened near the injection site over time (Fig 7D & 7E). These findings demonstrate that the phase separation of WNK1 in vivo is due to changes in cytoplasmic crowding rather than cell size. These findings implicate WNK kinases as intracellular molecular crowding sensors that utilize phase separation as a mechanism to control cell volume.

## Discussion

WNK kinases have long been appreciated to participate in cellular osmosensing, but the mechanisms by which they activate downstream signaling following volume contraction have been obscure. In this work, we show that WNK1 responds to hyperosmotic stress by detecting a crowded cytosol. It does so by undergoing LLPS, forming functional condensates that activate the WNK-SPAK/OSR1 pathway to trigger downstream NKCC1/KCC phosphorylation, net ion influx, and RVI (Fig 8). WNK1 phase behavior occurs within physiological constraints and is mediated by its large CTD, which is highly conserved with respect to its size, regional disorder tendency, and domain organization despite poor sequence identity across evolution. These features allow even the most ancient WNKs to quickly detect small changes in hypertonicity. Taken together, our results reveal an evolutionarily conserved function for WNK kinases as physiological crowding sensors, and identify WNK1 as a critical component of a unique type of signaling condensate that manages acute reductions in cell volume.

**Figure 8.**
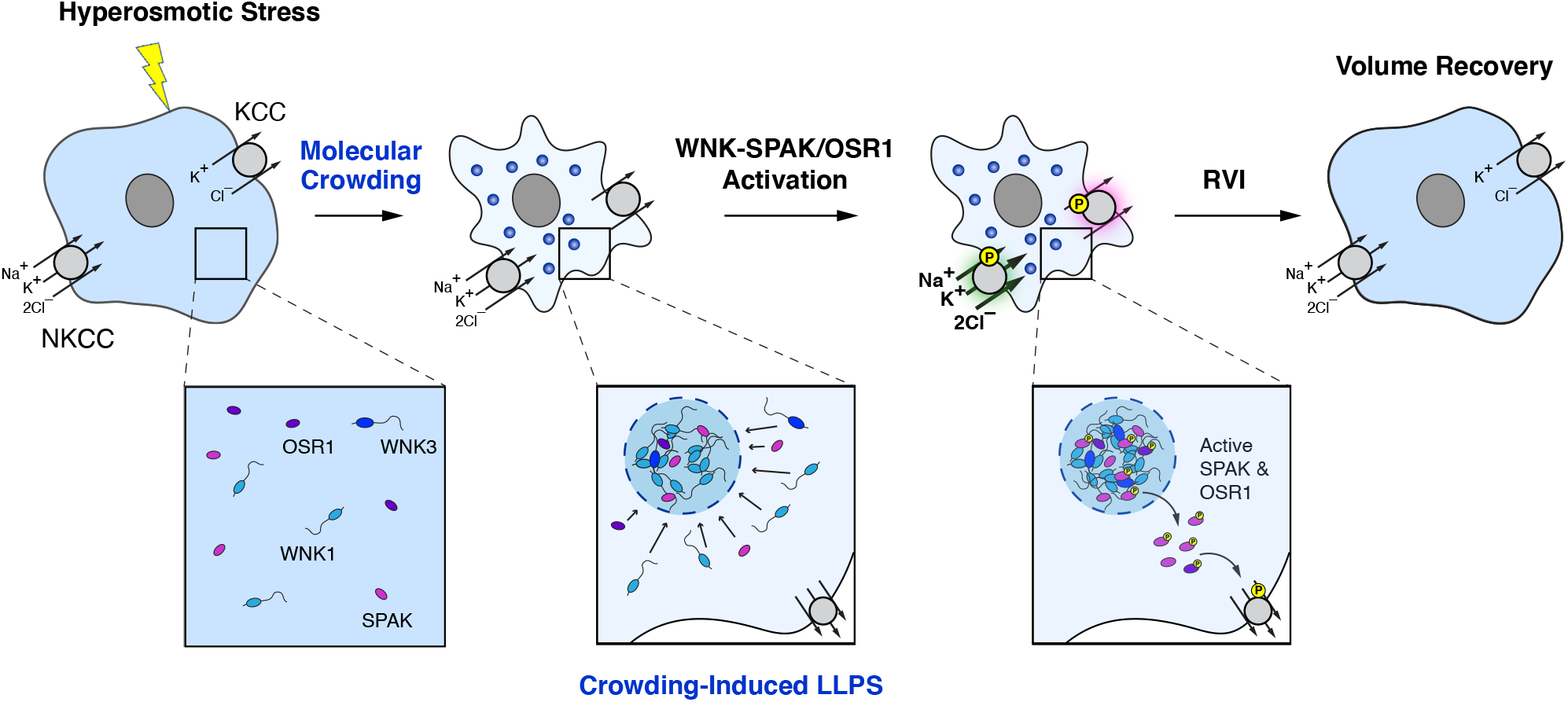
Model for WNK-SPAK/OSR1 pathway activation during hyperosmotic stress. Hyperosmotic stress causes cells to shrink, increasing the degree of macromolecular crowding. This triggers phase separation of WNK1 and other WNK-SPAK/OSR1 pathway components into condensates. Localization of the pathway within this confined space favors the phospho-activation of SPAK and OSR1, which exhibit increased cytosolic localization in their phosphorylated state. Material leaves the WNK1 condensates over time, allowing the kinases to directly phosphorylate NKCC1 and the KCCs at the plasma membrane, resulting in net ion influx and RVI.

The theory of macromolecular crowding as a cell volume signal and as a stimulus for cytoplasmic phase separation was proposed many years ago (Parker, 1993; Walter and Brooks, 1995; Zimmerman and Harrison, 1987). These concepts have been underappreciated; however, a paradigm shift in our understanding of cytoplasmic organization has prompted their reappraisal (Hyman et al., 2014). Hypertonicity diminishes solvation of the cytosol, creating an ideal milieu for interactions between multivalent proteins that are prone to phase behavior. This suggests that crowding-induced phase separation caused by hyperosmotic stress broadly modulates the activity of biomolecules that tend to form condensates (Jalihal et al., 2020).

The ability of WNK1 to activate in response to crowding addresses a longstanding paradox in the WNK signaling field. As shown here, the WNK-SPAK/OSR1-NKCC pathway is upregulated by hypertonicity; however, intracellular chloride and potassium depletion are also potent WNK activators (Piala et al., 2014; Pleinis et al., 2021). These observations have been difficult to reconcile, as hypertonic stress raises intracellular ionic strength due to exosmosis (O’Neill, 1999). Because this elevation in intracellular chloride and potassium concentrations should inhibit WNK kinase activity, the stimulatory effects of hypertonicity seem counterintuitive. Our results provide evidence that the crowding-dependent concentration of WNK kinases within condensates allows them to autoactivate while bypassing the inhibitory effects of high ionic strength caused by cell water loss.

WNK kinases harbor a very large intrinsically disordered C-terminal domain, usually greater than 100 kilodaltons in mass, whose function has been incompletely understood. Our studies show that the entire domain is critical for WNK1 to undergo phase separation in response to modest reductions in cell volume. The WNK CTD can therefore be thought of as a physiological crowding sensing domain. Though the sequence identities of WNK CTDs vary widely across evolution, their regional features in terms of disorder, low complexity, and prion-like signatures share striking similarities. In mammalian WNK1, regions at both the proximal and distal ends of the CTD are particularly prone to phase behavior, and coiled-coil domains within these regions mediate condensate coarsening, augmenting kinase retention within the condensed phase to optimize downstream SPAK/OSR1 activation. Moreover, deletion of the distal end of the CTD impairs condensate formation, reducing the sensitivity of WNK1 to crowding. Analysis of the *Drosophila* WNK CTD suggests that these features are evolutionarily conserved. Thus, it appears that the large size of the CTD is a conserved feature that allows WNK kinases to react to physiologic changes in cytosolic crowding. These findings have implications for understanding how IDRs mediate physiological sensing functions in vivo.

Though WNK1 phase behavior is essential for its hypertonic activation, other mechanisms appear to be involved. For example, the cell swelling-activated kinase ASK3 forms inactive condensates during hyperosmotic stress (Watanabe et al., 2021). Since ASK3 is a WNK1 suppressor (Naguro et al., 2012), its deactivation within condensates likely augments WNK1 activity during crowding. In addition, the swell-activated chloride channel LRRC8A exhibits physiological activity during hypertonic stress, as ablation of channel activity during hypertonicity aberrantly increases [Cl]_i_ and exacerbates cell shrinkage due to impaired WNK1 activity (Serra et al., 2021). These mechanisms likely serve permissive roles that mitigate WNK1 inactivation during shrinkage (Goldsmith and Huang, 2021). A third study found that the purified WNK1 and WNK3 kinase domains activate in vitro in the presence of molecular crowders (Akella et al., 2021). This effect was associated with dehydration of the kinase domain, suggesting that solvent effects participate in a direct osmosensing mechanism. This finding may be relevant to the mechanism of condensate-mediated WNK1 autoactivation, since condensed phases provide alternate solvent environments (Nott et al., 2015). Conceivably, reduced WNK1 hydration status within condensates may trigger conformational changes within the kinase domain that help it to override the inhibitory effects of ionic strength.

The findings reported here have broad implications for WNK-dependent signaling in health and disease, as the mechanisms that mediate RVI have been leveraged by higher organisms to control advanced physiologic processes. For example, mutations in the WNK signaling pathway cause hypertension and hyperkalemia due to the overactivation of NCC, an NKCC1-like SLC12 salt cotransporter localized to kidney distal convoluted tubule (DCT) (Subramanya and Ellison, 2014). Within this nephron segment, WNK activation is associated with the formation of DCT-specific biomolecular condensates called WNK bodies (Boyd-Shiwarski et al., 2018). The assembly of these membraneless structures is driven by KS-WNK1, a truncated WNK1 isoform that retains the entire disordered CTD shown here to be critical for phase behavior. This suggests that KS-WNK1 scaffolds WNK-dependent phase transitions that participate in volume and K+ homeostasis. NKCC1-dependent transport contributes to colonic epithelial K+ secretion (Nickerson and Rajendran, 2021), choroid plexus function (Steffensen et al., 2018), and ischemic stroke (Bhuiyan et al., 2016); these processes likely require WNK phase behavior. Furthermore, drugs targeting the WNK-SPAK/OSR1 pathway may influence condensate dynamics. ZT-1a, a potent small molecule SPAK inhibitor, disrupts interactions with WNK1 and mitigates ischemic injury following stroke (Zhang et al., 2020). It would be of interest to evaluate whether ZT-1a modulates condensate-dependent WNK signaling, as this may have implications for future therapeutics for the WNK-SPAK/OSR1 pathway, and for condensate drug design in general.

Phase separation has emerged as a fundamental property of biomolecules that is reshaping theories regarding cytoplasmic and nuclear organization. However, protein phase behavior is often only loosely connected to physiological relevance (Leslie, 2021; McSwiggen et al., 2019). Our results provide evidence that phase separation of the WNK signaling pathway is functionally relevant, occurring at native protein concentrations and at levels of osmotic stress that are within a physiological range commonly experienced during life. Many of the biophysical principles regarding the inherent molecular features of phase separating systems are evolutionarily encoded within WNK kinases to link condensate formation to cell volume regulation. Thus, the results presented here provide strong support for the relevance of biomolecular condensates in cellular physiology.

## Methods

### Plasmid Construction

See Table S2 for a list of all plasmid constructs used in this study. All mammalian WNK1 clones used for this study were derived from the original untagged rat WNK1 cDNA (AAF74258.1), which encodes a 2126 amino acid kinase-active isoform isolated from brain (Xu et al., 2000). This cDNA was modified to correct a variant serine present in residue 2120 of the original cDNA back to a conserved glycine to match the canonical Uniprot sequence Q9JIH7-1, and subcloned into the *EcoRI* and *XbaI* sites of pcDNA3.1. In the plasmids table, this complete CDS is referred to as rWNK1-S-G, but is referred to as “full-length WNK1” in this study. C-terminal eGFP-tagged WNK1 was reported previously (Boyd-Shiwarski et al., 2018). To generate N-terminal mRuby2-tagged WNK1, a gBlock encoding mRuby2 fused to the WNK1 N-terminus via a GRGS linker was synthesized (IDT) and swapped with an *EcoRI*/*NotI* N-terminal fragment in WNK1. Truncated N-terminal mRb2 tagged 1-494 and 1-1242 fragments constructs were generated by PCR off the full-length template and ligation to linearized pcDNA3.1. To generate the double coiled-coil mutant, two gBlocks encoding proline substitutions in the mid and distal coiled coil domains were swapped in stepwise fashion with corresponding regions into the full length mRuby2-WNK1 construct in pcDNA3.1 via restriction cloning. For the Cry2 plasmids, WNK1 fragments were generated by PCR or as synthetic gBlocks and were ligated via restriction cloning to the *XmaI* site of linearized Cry2PHR or Cry2Olig (E490G) vectors (Addgene #26866 and #60032), positioned in-frame between the PHR domain and mCherry.

For the dmWNK constructs, a gBlock encoding *Drosophila* codon-optimized human kinase-dead SPAK (HsSPAK^D210A^) was synthesized and inserted into the multicistronic vector pAc5 STABLE2 Neo, which contains a GFP cassette (Gonzalez et al., 2011) (Addgene #32426), by Gibson assembly to generate pAc5-GFP:T2A:HsSPAK^D210A^. dmWNK fragments were then PCR amplified from a full-length clone (Wu et al., 2014) and inserted into this vector in-frame with the GFP cassette by Gibson assembly to generate a series of N-terminal GFP-tagged dmWNK truncation constructs. To delete WNK1 from HEK-293 cells, we used a previously reported bicistronic PX330 plasmid (Addgene #42230) harboring the Cas9 CDS and a single guide RNA (sgRNA) targeting exon 1 of WNK1 (Roy et al., 2015b).

To generate the Cas9/WNK3-sgRNA plasmid, a 20bp guide sequence (WNK3.5) targeting exon 1 of human WNK3 was ligated to the *BbsI* site of PX330 (PX330-WNK3.5). To confirm WNK3 sgRNA efficacy, a complementary pair of 30bp oligonucleotides spanning the WNK3.5 sgRNA target cut site were ligated to a pHRS reporter plasmid (PNABio) (Pereira et al., 2016). To generate mNeonGreen WNK1 knock-in cells, a 20bp guide sequence targeting the 5’ end of WNK1 exon 1 was ligated to the *BbsI* site of PX459 (Addgene #62988). In addition, we generated a plasmid harboring a homology-directed repair knock-in template by inserting the mNeonGreen CDS and 5’ and 3’ homology arms flanking the WNK1 insertion site into pUC18 via overlap extension PCR and Gibson assembly. The homology arms were amplified from extracted HEK-293 cell genomic DNA (QuickExtract DNA extraction solution, Lucigen). The fidelity of all constructs was confirmed by Sanger sequencing prior to use.

### Cell culture and transfection

HEK-293 and U2OS cells were cultured in a 37-degree 5% CO_2_ incubator in complete media (glucose-supplemented DMEM with 10% FBS, L-glutamine, and penicillin/streptomycin). Plasmids were transfected using Lipofectamine 3000 (ThermoFisher) per manufacturer instructions. If transfected cells were used for imaging, they were re-plated on coverslips or MatTek dishes 24h post-transfection. GBM43 cells were cultured in a 37-degree CO2 incubator in Neurobasal media supplemented with B27-A and N2 (ThermoFisher), basic FGF, EGF, L-glutamine, and penicillin/streptomycin *Drosophila* S2-Gal4 cells (provided by Dr. Adrian Rothenfluh) (Acevedo et al., 2015) were maintained at 26°C, in Schneider’s Drosophila media supplemented with 10% Fetal Bovine Serum (ThermoFisher Cat. No 21720024). Drosophila constructs were transfected with plasmid DNA using TransIT®-Insect transfection reagent (Mirus Bio).

### Gene edited cell lines

WNK1/WNK3 double knockout (DKO) HEK-293 cells were generated via methods similar to (Roy et al., 2015b). Cells were grown in 6-well dishes, cotransfected with WNK1-PX330 plasmid and pEGFP-N1, used as a fluorescent marker for sorting transfected cells. 48h later, cells were harvested in PBS + 2% FBS, dilutely sorted into 96-well plates using a FACSAria II cell sorter (BD Biosciences), and clonally expanded. Knockout clones were identified by immunoblotting with validated WNK1 antibodies (Figure S2 and (Boyd-Shiwarski et al., 2018)). To knock out WNK3, the WNK3.5 sgRNA was first functionally validated using a surrogate reporter method (Pereira et al., 2016; Ramakrishna et al., 2014). PX330-WNK3.5 was cotransfected into HEK-293 cells with pHRS-WNK3.5, a reporter plasmid that harbors an out-of-frame GFP cassette 3’ to the target cut site. On-plasmid CRISPR-mediated nonhomologous end-joining (NHEJ) shifts the GFP CDS in-frame, resulting in green fluorescence, which was confirmed for cells expressing the WNK3.5 sgRNA by epifluorescence microscopy 48h post transfection. The PX330-WNK3.5 plasmid was then used to knock out WNK3 from WNK1 KO cells using methods identical to those described for WNK1 deletion.

To generate endogenous mNG-tagged WNK1 cells, PX459-WNK1, encoding Cas9 and WNK1- sgRNA, was cotransfected with the pUC18-WNK1/mNeonGreen homology directed repair template plasmid. 72h post-transfection, cells were sorted by FACS for mNeonGreen fluorescence, plated dilutely into 96-well plates, and clonally expanded. Individual clones were validated by immunoblotting for WNK1 and the mNG tag, and by amplification of the target site across the 5’ and 3’ homology arms by genomic PCR, TA cloning into pGEM-Teasy (Promega), and Sanger sequencing.

### Antibodies

To generate the guinea pig total NKCC1 antibody, we targeted amino acids 929-999 of mouse NKCC1 (NCBI NP_033220.2; CCDS 37826.1). This epitope is contained within a unique loop of the NKCC1 C-terminus, and is distinct from sequences in other SLC12 cotransporters, including the closely related NKCC2 (*SLC12A1*) (Carmosino et al., 2008). A fusion protein encoding the NKCC1 epitope fused to GST was expressed by IPTG induction in BL21 E.coli, purified using a HiTrap glutathione S-Transferase column, and eluted with 10mM reduced glutathione. The free glutathione was removed by overnight dialysis and purity was confirmed by SDS-PAGE. Antibody production in guinea pigs was carried out by Pocono Rabbit Farm and Laboratory (Canadensis, PA). Validation of the antibody in global NKCC1 KO mice (Flagella et al., 1999) is shown in Figure S1A. We were unable to detect phosphorylated NKCC1 using several commercial antibodies; therefore, we used a previously validated antibody that was directed to phosphorylated Threonine-58 of NCC (*SLC12A3*) (Sorensen et al., 2013). The epitope recognized by this antibody (amino acids 54-62 of mouse NCC; FGHYNpTIDVV) cross-reacts with a phospho-activation site present in mouse NKCC2 and NKCC1 (Gimenez and Forbush, 2003; Lee et al., 2013). Phosphorylation at this site is a signature of NKCC activation (Flemmer et al., 2002); thus, the pT58 antibody was used to provide a readout of activated NKCC1 in HEK-293 cells. TIAR was detected using a previously reported antibody (Daigle et al., 2016). Table S3 provides a complete list of antibodies that were used in the study.

### Live cell imaging of WNK condensates

HEK-293 cells were plated in 6-well dishes in complete media and allowed to grow to 80-90% confluency. Cells were transfected in antibiotic-free media the following day with standardized amounts of mRuby2 tagged WNK1 plasmids. Cells were re-plated 24h post-transfection on poly-D-lysine coated MatTek dishes. Tonic solutions and media were prepared and warmed to 37°C on the day of imaging. Cell media was replaced with Leibovitz L-15 media and dish was loaded into a climate-controlled stage set to 38°C of a Leica SP8 live cell resonant scanning confocal microscope using a Leica HC Pl APO CS2 63X, 1.40 numerical aperture oil objective. Scan speed was set to 1000 Hz and images were captured in a 2048x2048 resolution format. Cells were screened and selected for imaging with similar mRuby2 signal intensities to evaluate condensate behavior in cells with relatively equal amounts of WNK1 protein by taking an intensity measurement in Fiji prior to imaging. Stress media (50 mM sorbitol) was added after the second stack was taken to establish a pre-stress baseline. Confocal stacks (10-18 steps, 1µm step size) capturing mRuby2 signal were taken every 15 seconds for 10 minutes.

Spinning disk confocal fluorescence live cell imaging was performed on a Nikon Ti-E-2000 inverted microscope equipped with an Andor Revolution XDi spinning disk, a 100X objective (Nikon CFI Plan Apo TIRF Lambda, 1.49 NA), Piezo XYZ stage (Nikon), iXon 897 Ultra back-illuminated camera (Andor Technology), a laser combiner (Andor Technology) containing 405, 488, 515, 568, and 647 nm excitation capabilities, Perfect Focus system (Nikon), a Dell 5400 Workstation with IQ2 imaging software (Andor Technology), and an active isolation air table (TMC) (Shiwarski et al., 2019). The system was surrounded by a full heated enclosure to maintain physiological temperature and humidity. eGFP and mRuby WNK1 expressing cells were grown on coverslips and transferred to a metal imaging chamber (AttoFlour, Invitrogen) in Leibovitz’s L- 15 media + 1% FBS. eGFP and mRuby were imaged using 488 nm laser excitation with 525/50 nm emission filter, and 561 nm laser excitation with 620/60 nm emission filter, respectively. Exposure duration, laser power, camera gain, and Z-step sizes were optimized to achieve fast 3D imaging at Nyquist resolution of WNK1 condensate formation while minimizing photobleaching during the time-lapse imaging. 3D rendering of the image stacks were performed using the IQ2 built-in animation features and Imaris software (Bitplane).

For live cell imaging in *Drosophila* S2-Gal4 cells, 48 hours following transfection, cells expressing GFP-dmWNK constructs were resuspended and transferred in 500 µl aliquots to 1.5mL centrifuge tubes. The cell suspension was then diluted stepwise with water in 10% increments over 20 minutes to a final concentration of 60% cell media to 40% water (174 mOsm). 100 µl of the diluted cell/media mixture was then transferred to 8-well live cell imaging plates (Ibidi, Cat. No. 80827), mixed with water (hypotonic condition, 87mOsm, to completely dissolve condensates) or sorbitol (Fisher Scientific, Cat. No. BP439-500) (hypertonic condition, 376 mOsm, final sorbitol concentration 250 mM) and allowed to equilibrate for 30 minutes. In some experiments, sorbitol was added to achieve varying concentrations. Osmolality was measured using a vapor pressure osmometer (Vapro 5520, Wescor). Cells were imaged using an Olympus CKX53 inverted fluorescence microscope with excitation between 488 nm and 510 nm using the 40X objective and representative images were captured. The GFP channel was isolated and converted to inverted grayscale images in Fiji. For puncta counting experiments, the fraction (percentage) of cells with condensates was quantified by counting the number of cells with condensates and dividing by total number of cells at varying osmolalities. Movies of transfected S2 cells were acquired with a Leica SP8 with 405, 488, 561, and 633 nm laser lines, using a HC PL APO CS2 63x, 1.4 numerical aperture oil objective with a 1x zoom factor. Images were acquired in an 8-bit 1024x1024 resolution format with 1 frame average and 2 line averages while scanning at 200 Hz.

### Immunofluorescence confocal microscopy in fixed cells

Cells were plated on Biocoat coverslips and allowed to adhere overnight. Hypertonic stress media and fixative (4% paraformaldehyde in hypertonic stress media) solutions were warmed to 37° C. Media was aspirated off and hypertonic stress media was applied to cells. Cells were incubated in stress media in a 37°C CO_2_ incubator for 5 minutes. The stress media was aspirated and warmed hypertonic fixative was applied to cells for 15 minutes at 37° C. Following fixation, the paraformaldehyde was quenched with 50mM NH4Cl, and the cells were permeabilized with 0.1% Triton X-100. Following washing with PBS w/ Ca^2+^ and Mg^2+^ (PBS^+/+^) the cells were incubated for 1h at room temperature with blocking solution containing FBS and fish gelatin (1% w/vol). Primary antibody was diluted in block solution and applied overnight at 4°C. The next day, following washes with block solution, fluorescent secondary antibodies (diluted in block solution) were applied in the dark for 2h at room temperature. Following additional washes in block solution, cells were stained with a To-Pro3 in block solution (diluted 1:5000; Invitrogen, T3605) at room temperature for 20 minutes. After a final set of washes, coverslips were mounted onto slides using ProLong Glass (Invitrogen, P36980).

For standard confocal microscopy, images were acquired with a Leica TCS SP5 CW-STED with 488, 552, and 633 nm laser lines, using a HCX PL APO CS 40x, 1.25 numerical aperture oil objective with a 4x zoom factor. Images were acquired in a 12-bit 1024x1024 resolution format with 4 frame averages and 8 line averages. Crosstalk between channels was mitigated with sequential channel scanning. Confocal images with high-resolution adaptive deconvolution processing were obtained with a Leica Stellaris 8 FALCON FLIM white light laser scanning confocal microscope, using a Leica HC Pl APO CS2 63X, 1.40 numerical aperture oil objective with a 3x zoom factor. mNeonGreen and mRuby2-specific spectra were imported to LAS-X from FPBase. Scan speed was set to 400Hz and images were captured in a 2048x2048 resolution 16- bit format with 4 line averages. Confocal images were processed adaptively using Lightning deconvolution software (Leica). Sequential scanning was used to prevent channel crosstalk. In all cases, multiple confocal fields were obtained from at least two biological replicates, obtained over the course of 2-3 independent experiments.

### CLEM

HEK293 cells were grown in 6-well dishes and transfected with full-length mRuby2-WNK1. 24h post-transfection, cells were re-plated onto poly-D-lysine-coated MatTek dishes and allowed to grow for an additional 24h. Growth media was then replaced with complete media supplemented with 500mM sorbitol (hypertonic). All cells were incubated in a 37-degree CO_2_ incubator for 3 minutes before applying fixative (4% paraformaldehyde in isotonic or hypertonic media, respectively) at 37°C for 1h. Fixed cells were washed twice with PBS at room temperature without agitation and stored in PBS at 4°C until imaged.

Fluorescent images were taken on a Nikon A1 confocal microscope. Samples were first imaged with a 0.50NA 10x objective using the “ND Acquisition + Large Images” addon to create a stitched montage of images of the entire coverslip, exciting with a 561nm laser and collecting through a 595/50 filter. Regions of interest were reimaged with a 1.40NA 60x objective using Nyquist sampling (0.21µm/px) and a 3x3 stitched large image in a 20µm z-stack with 0.17µm steps. A straight scratch through the sample made with a pipette tip was used as a fiduciary mark for later alignment with the SEM data.

Following high-resolution confocal field-of-view imaging, the samples underwent post-fixation and en bloc heavy metal staining to prepare for the SEM backscatter imaging. The samples were first placed in a 1% OsO4 solution for 20 minutes, followed by three 5-minute PBS washes. Next, the samples were then placed in a 1% Tannic Acid solution for 10 minutes, followed by three 5-minute ddH_2_O washes. Finally, the samples were then placed in a second 1% OsO4 solution for 15 minutes followed by three 5-minute ddH_2_O washes. Samples were then stained en bloc with aqueous 2% uranyl acetate for two hours at room temperature followed by three 5-minute ddH_2_O washes. The final en bloc stain used was a lead aspartate solution placed on the samples and incubated at 60°C for one hour followed by three 5-minute ddH_2_O washes. After post-fixation and en bloc staining, the samples were be dehydrated and embedded in resin. Samples were dehydrated with a graded series of ethanol; 30%, 50%, 70%, 90%, 100%, followed by infiltration with four 1-hour changes of EPON embedding resin (Franks et al., 2017). Pre-formed resin blocks that fit the SEM stage are pressed into the unpolymerized resin of the samples and allowed to polymerize at 37°C for 24 hours and then 65°C for 48 hours. Once the samples were acclimated to room temperature, the coverslip from the MatTek cell dish was removed with a freeze/thaw method, alternating the dish briefly sitting in liquid N_2_ and 100°C H_2_O until the glass coverslip was removable.

The sample block was then transferred to the SEM holder and polished with a JEOL Handy Lap with a ThorLabs 0.3µm Calcined Alumina Lapping Sheet. Roughly 20 strokes of the Handy Lap were adequate to remove the outer membrane of the embedded cells. The sample was then coated in carbon and placed in the JEOL JSM-7800 SEM equipped with a backscatter detector for imaging. Samples were SEM imaged at 7kV. A 250x montage of the entire face of the block was taken in order to align with the 10x fluorescent image montage. The polishing, carbon coating, and 250x SEM imaging was repeated until fiduciary marks could be identified and the precise location on both the SEM and fluorescent data could be established and correlated. Once the areas where the 60x confocal fluorescent images were taken and identified, SEM montage of the regions of interest were taken at 10Kx or 15Kx. Polishing, carbon-coating, and 10Kx or 15Kx SEM montage imaging was also repeated until cellular level fiduciary marks could be identified within the Z-planes of the fluorescent images and aligned. The high magnification SEM montages were stitched using Nikon Elements software and the Fluorescent and EM data were aligned.

### FRAP

Fluorescence recovery after photobleaching (FRAP) studies were performed during live confocal fluorescence microscopy as previously reported (Boyd-Shiwarski et al., 2018) using the imaging system described above. Images were acquired with 100X objective (Nikon CFI Plan Apo TIRF Lambda, 1.49 NA) every 300 ms for 1 min. FRAP of eGFP-L-WNK1 condensates was performed using an Andor FRAPPA unit with the 488 nm laser at 100% power, 100 µs dwell time and two repeats for each spot. Image series were imported into FIJI ImageJ for analysis. To calculate the percent recovery after photobleaching we normalized the bleached region to a control non-bleached region over time. The initial fluorescence intensity of the condensate prior to photobleaching was averaged and used to normalize the data. To isolate the recovery of the mobile fraction, the minimum fluorescence intensity for each region was subtracted from the normalized fluorescence intensity over time, and normalized to constrain the prebleach signal intensity to 100% and the post bleach intensity to 0%. Data was plotted as percent recovery vs. time in seconds using Prism 9 (GraphPad).

### Optogenetic screen for phase separating WNK1 domains

All optogenetic live-cell imaging experiments were performed on a Nikon A1 laser-scanning confocal microscope system outfitted with a Tokai HIT stagetop incubator utilizing 40X and/or 60X oil immersion objectives (CFI Plan Apo Lambda 60X Oil, Nikon). Following transfections and/or treatments, medium was changed to phenol red-free growth medium (GIBCO) and cells were allowed to equilibrate on the preheated (37°C and 5% CO2) stagetop incubator for 10 min prior to imaging. Acute blue light stimulation was achieved by utilizing the 488nm laser line and the stimulation module within Nikon Elements imaging software. Activation duration was set to 1s and laser power was set at 20%. Stimulation regions of interest (ROIs) were drawn over fields of view prior to image acquisition. Following 2-5 baseline images, laser stimulation was performed and cells were imaged for up to 10 min post-activation. Timing and order of image acquisition was alternated across experiments between experimental groups. Data presented are representative of at least two independent experiments utilizing three or more biological replicates per experiment.

### In-cell phase diagrams

Cells were transiently transfected with mRb2-tagged WNK1 constructs. 24h later, the cells were re-plated on poly-D-lysine coated MatTek dishes. The following day (∼48h post transfection), media was replaced with 1mL pre-warmed Leibovitz L-15 media. Dishes were mounted into an environmental stage set to 38°C on an inverted Leica SP8 confocal microscope with a HC PL APO CS2 63X, 1.40 numerical aperture oil objective. 6x6 tile scans were obtained every 60 seconds for 10 minutes. The scan speed was set to 700 Hz with resonant scanning deactivated, and images were acquired in a 2048x2048 resolution format. Fixed laser power for the 552 nm laser line was set using the 1-494 mRuby2 tagged WNK1 construct as this was the most highly expressed construct, and thus the brightest. This was done to minimize oversaturation in each set of transfected cells. Stress media was added after a pre-stress tile scan was completed to obtain baseline fluorescence intensities. The stresses were composed of Leibovitz media with varying amounts of sorbitol, ranging from 50 mM (350mOsm) to 250 mM (550mOsm).

In each of the tile images, mRuby2-fluorescent cells that were clearly captured within the field of view for the duration of the stress time course were identified, assigned an identification number for blinded evaluation (Fig S4A), and pre-stress intensities were measured in FIJI. The blinded image sets were categorized by stress condition and transfected WNK1 construct but were deidentified via a randomized coding system prior to passing them on to three independent screeners. The screeners scored the degree of phase separation (no phase separation, nucleation, and spinodal decomposition) for each numbered cell. Score discrepancies were re-evaluated by the screeners as a group on a case-by-case basis to arrive at consensus. Following unblinding, data points were plotted on a graph with protein concentration on the X-axis, and osmolality on the Y-axis to compose phase diagrams. Binodal phase boundaries were drawn at the level of pre-stress fluorescence intensity where the WNK1 constructs uniformly adopted a nucleated morphology following exposure to osmotic stress, and spinodal phase boundaries were drawn in similar fashion for constructs that underwent post-stress spinodal decomposition.

### Preparation of cell and tissue lysates and immunoblot analysis

To examine the natively expressed WNK-SPAK/OSR1 pathway under hyperosmotic stress, unedited HEK-293 cells or endogenous mNG-tagged WNK1 cells were grown in 6-well dishes. Once cells reached 90% confluence (∼24h), growth media (complete) was replaced with fresh complete media supplemented with various amounts of sorbitol. Following the application of isotonic or stress media, cells were incubated in a 37-degree CO_2_ incubator for 5 minutes, and then harvested (∼5 minutes) in the same applied isotonic or stress media. Collected samples were immediately pelleted at 4°C for 5 minutes and the supernatant discarded, totaling 15 minutes of exposure to osmotic stress. For experiments with transfected WNK1 constructs, WNK1/WNK3 DKO cells were grown in 6-well plates and transfected, in duplicate, with varying amounts (0.5- 3.5ng) of one of the following mRuby2-tagged WNK1 constructs: full-length,1-494 fragment, 1- 1242 fragment, or the double coiled-coil mutant. Based on pilot studies, transfection amounts were adjusted to minimize differences in protein expression among each of the constructs. 24h post-transfection, cells were re-plated, dividing each well of transfected cells in two. After an additional 24h, growth media (complete) from one divided well was replaced with fresh complete media (iso), and growth media from the second divided well replaced with complete media supplemented with 50mM sorbitol (hyper). Cells were placed in a 37-degree CO_2_ incubator for 20 minutes and then harvested in the same applied media (∼5 minutes). Collected samples were immediately pelleted at 4°C for 5 minutes and the supernatant discarded, totaling 30 minutes of exposure to osmotic stress. All other unedited cells and candidate clones were washed with chilled DPBS and collected in chilled PBS supplemented with protease (Thermo Fisher Scientific) and phosphatase (Roche) inhibitor cocktails. Collected samples were immediately pelleted at 4°C for 5 minutes.

Cytosolic proteins were extracted from cell pellets with ice-cold Detergent Lysis Buffer (50mM Tris-Hcl pH 8.0, 1% IGEPAL CA 630, 0.4% sodium deoxycholate, 62.5mM EDTA, supplemented with protease and phosphatase inhibitors). Lysates were incubated on ice for 20 minutes before pelleting out insoluble materials via hard centrifugation at 4°C for 10 minutes. Supernatant protein concentrations were determined using the Pierce BCA Protein Assay Kit (Thermo Fisher Scientific). Equal amounts of protein (20ug) were denatured in Laemmli buffer, maintained at room temperature for 10 minutes, and loaded onto 4-20% Criterion TGX precast gels (Bio-Rad) alongside Precision Plus All Blue Standard (Bio-Rad) for SDS-PAGE. Once separated, the proteins were transferred to a nitrocellulose membrane using Bio-Rad Trans-Blot Turbo for immunoblotting. The resulting signal was visualized with enhanced chemiluminescence and measured using a Bio-Rad ChemiDoc imager.

For immunoblotting data obtained in transfected osmotically stressed cells, gel lanes represent individual biological replicates, and are representative of data obtained over three to five independent experiments, which were used for quantification and statistical analysis.

For protein isolation from tissue, whole kidney and brain cortex was homogenized in ice-cold RIPA extraction buffer (Thermo Scientific) with freshly added protease and phosphatase inhibitors. Following agitation at 4°C for 30min, homogenates were centrifuged and supernatants were collected. Equal amounts of protein, as determined by BCA protein assay, were fractionated by SDS-PAGE on 4-15% Criterion TGX precast gels, transferred to nitrocellulose, and immunoblotted with NKCC1 antibody, as described above. ECL signal was visualized by autoradiography.

### Cell volume measurements

HEK-293 WNK1/WNK3 DKO cells were plated in 6-well dishes in complete media and allowed to grow to 80-90% confluency. Cells were transfected in antibiotic-free media the following day with standardized amounts of mRuby2 tagged WNK1 plasmids and pEGFP-N1. Since eGFP distributes uniformly throughout the nucleus and cytoplasm, it was used as a fluorescent marker for transfected cell volume. Cells were re-plated 24h post-transfection on poly-D-lysine coated MatTek dishes. Tonic solutions and media were prepared and warmed to 37°C on the day of imaging. Cell media was replaced with Leibovitz L-15 media supplemented with 10 µM Cariporide, with or without 20 µM Bumetanide. Both cariporide and bumetanide were solubilized in DMSO; the addition of these inhibitors increased osmolality by 7 and 13mOsm, respectively. Following a 10 min preincubation period, the plated cells were imaged with a Leica SP8 live cell resonant scanning confocal microscope on a climate-controlled stage set to 38°C. Confocal stacks (20-30 steps, 2µm step size) capturing whole-cell eGFP signal were taken with a Leica 40x, 1.1 numerical aperture water immersion objective every 30 seconds for 20 minutes at a scan speed of 8000Hz. For each experimental condition, a control set of calibration measurements was taken to account for natural drift in cell volume in Leibowitz media on the heated stage over a 20-minute time course. To evaluate volume changes after hypertonic stress, following the first 60 seconds of imaging, sorbitol was added to achieve a final concentration of 50mM, which increased solution osmolality by 50mOsm. Confocal stacks were rendered in the LasX 3D software platform. Uncommonly, field boundaries were cropped during post-processing to exclude floating cells that would confound volume calculations. Following background subtraction, Otsu thresholding was applied to obtain binary images. The integrated binary whole-cell GFP signal from each confocal stack was used to calculate the percent cell volume per imaging field; this reliably reflected volumetric changes over the time course.

For cell volume calculations, percent volume per field measurements were normalized to 100% at time zero. Changes in cell volume over time were adjusted by the control set of drift calibration measurements. RVI was defined as the percent volume recovery during the time course following exposure to hypertonic stress. To account for variability in baseline volume reduction after hypertonic exposure, this percentage was expressed relative to the volume lost, using the following equation:

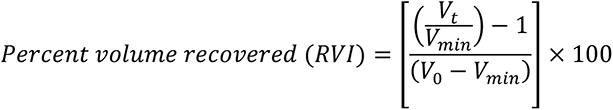

Where V_0_ = the volume at time zero (normalized to 1), V_min_ = the minimal fractional cell volume measured immediately following exposure to hypertonicity, and V_t_ = the fractional cell volume at a given time point during the RVI phase.

### Rubidium influx assay

WNK1/3 DKO cells expressing full-length or 1-494 WNK1 were cultured in PDL coated 24-well plates with DMEM culture media until 90% confluent. The culture medium was removed and cells were rinsed with an isotonic buffer (310 mOsm, containing 134 mM NaCl, 2 mM CaCl_2_, 0.8 mM NaH_2_PO_4_, 5 mM glucose, 25 mM HEPES and 1.66 mM MgSO_4_) as described before (Luo et al., 2020). To measure basal K^+^ (Rb^+^) influx, cells were exposed to the isotonic buffer (310 mOsm, pH 7.4, containing 5.36 mM Rb^+^,) in the absence or presence of NKCC1 inhibitor bumetanide (BMT, 10 µM), for 10 min at 37°C. To measure changes of Rb^+^ influx in HEK293 cells in response to hypertonic stress, cells were exposed to the hypertonic solution (370 mOsm adjusted by adding 60 mM sucrose, pH 7.4, 5.36 mM Rb^+^) for 5 min at 37°C. To terminate Rb^+^ influx, cells were washed with the isotonic or hypertonic washing solutions (Rb^+^ free) and lysed with 200 mL 0.15% SDS (room temperature) to release the intracellular Rb^+^. The intracellular Rb^+^ concentration in cell lysates was measured using an automated atomic absorption spectrophotometer (Ion Channel Reader, ICR-8000; Aurora Biomed, Vancouver, Canada). Total protein of cell lysates was measured by BCA assay. Rb^+^ influx rate was calculated and expressed as mg Rb^+^/mg protein/min. NKCC1-mediated Rb^+^ influx was determined by subtracting Rb^+^ influx value in the presence of bumetanide from the total Rb^+^ influx. To calculate shrinkage-induced bumetanide-sensitive Rb+ flux, we subtracted the baseline bumetanide-sensitive Rb+ transport measured under isotonic conditions (310mOsm), from the increase in bumetanide-sensitive Rb+ transport measured under hypertonic conditions (370mOsm).

### Crowding agent microinjection studies

Borosilicate glass filaments (O.D. 1.2 mm, I.D. 0.94 mm, 10cm length, Catalog #: BF120-94-10 Sutter Instrument) were pulled into micropipettes using a Sutter Instrument P-1000. A potassium acetate HEPES buffer (KH buffer, 125 mM K+ Acetate, 25 mM HEPES, pH 7.4), was prepared as the vehicle for microinjection and 1.5 µL was backfilled into the pulled micropipettes (Shiwarski et al., 2017). For isosmotic injections, stock KH buffer was injected into mNG-WNK1 cells using an Eppendorf FemtoJet (pi = 35 hPa, ti(s) = 0.2, pc(hPa) = 20) and InjectMan micromanipulators on a Nikon Ti-E-2000. Widefield fluorescence imaging and brightfield DIC was performed using a 100X objective (Nikon CFI Plan Apo TIRF Lambda, 1.49 NA), a GFP excitation/emission filter cube (Nikon), a stage top incubator (TokaiHIT), and a Prime 95B camera (Photometrics) run with Micro-Manager software (Edelstein et al., 2014). Time-lapse imaging (300 ms exposure, 30 s interval, for 15 min) was performed at 37°C prior to and during injection. A hyperosmotic solution was prepared by adding KCl (100 mM) 1:1 to the KH buffer to yield a 50 mM final concentration. To induce molecular crowding, a Ficoll solution (50% w/v Ficoll PM400, GE Healthcare, in 50 mM HEPES pH 7.4) was combined 1:1 with the KH buffer and microinjected into the cells as described for the isosmotic condition. In experiments injecting multiple solutions into the same cells, micropipettes were preloaded prior to the experiment and exchanged between treatments. Image analysis was performed using FIJI.

### Bioinformatic analysis

We used three algorithms to predict WNK protein disorder: SPOT-Disorder (Hanson et al., 2017), VSL2B (Obradovic et al., 2005), and VL3H (Peng et al., 2005). SEG was used to predict low complexity regions (LCRs) (Wootton, 1994). Short, medium, and long low-complexity segments were defined as having trigger window length (W), trigger complexity (K_2_(1)) and extension complexity (K_2_(2)) parameters of 12/2.2/2.5, 25/3.0/3.3, and 45/3.4/3.75, respectively. NCoils (Lupas et al., 1991) with a sliding window of 21 or 28 residues was used to predict coiled-coil domains. Marcoil (Delorenzi and Speed, 2002) was used to predict CC heptad registers in rat WNK1. Prion-like domains (PLDs) were predicted using PLAAC (Lancaster et al., 2014). A default core length of 60 and a relative alpha of 50% was used. Similar results were obtained when the relative background frequency alpha for *S. cerevesiae* was adjusted to 0% (i.e., 100% for the species being scored: human, rat, *Drosophila*, or *C. elegans*).

Details on the protein sequences used to analyze cross-species amino acid compositional bias in WNK CTDs are provided in Table S1. In most cases, UniprotKb was used to identify specific WNK kinases. Once the sequence for a putative WNK kinase was found, we confirmed its identity by verifying that the kinase domain contained the atypically placed catalytic lysine (Min et al., 2004) and chloride binding pocket that are defining features of the WNKs (Piala et al., 2014). The CTD was defined as the sequence downstream from the predicted PF2-like domain. Amino acid frequencies within the CTD were stratified using the Expasy ProtParam tool.

### Statistical analysis

Statistical details of experiments, including n, P-values, and specifics regarding post-hoc testing can be found in the figures, figure legends, and methods. Statistical analysis was performed using GraphPad Prism software and are presented as mean ± SEM. Comparisons between two groups were determined by Student’s t-test. Multiple comparisons were determined by one- or two-way ANOVA followed by post-hoc testing as indicated in the figure legends. Significance criterion was set at a P-value of ≤0.05. For colocalization studies in fixed cells, percent colocalization was determined by thresholded Mander’s Overlap Coefficients, using the JaCOP plugin in FIJI.

## Acknowledgments

This work was supported by National Institutes of Health grants K08DK118211 (C.B-S.); K99HL155777 (D.J.S.); R01DK098145 and R01DK119252 (A.R.S.); P30DK79307, S10OD021627, and S10OD028596 (Pittsburgh Center for Kidney Research); R01DK110358 (A.R.R.); and U.S Departments of Veterans Affairs grants I01BX002891 and IK6BX005647 (D.S.).

We thank Ora Weisz, Catherine Baty, Gerard Apodaca, and Donna Stolz for imaging support, Gabrielle Pittman and Katherine Querry for technical assistance, and Ossama Kashlan, Seth Childers, Jeff Brodsky, and Todd Lamitina for helpful discussions. This content is solely the responsibility of the authors and does not necessarily represent the official views of the U.S. Department of Veterans Affairs.

## Author Contributions

Conceptualization, C.B-S., D.J.S., A.R.S.; Methodology, C.B-S., D.J.S., C.J.D., D.S., A.R.R, A.R.S.; Investigation, C.B-S., D.J.S., S.E.G., R.T.B., D.E.M., L.N., J.M., J.W., W.T., E.N.A., J.F., M.C., K.A.C., C.J.W., C.C.W.; Resources, U.B.P., C.J.D., D.S., A.R.R., A.R.S. Writing – Original Draft, C.B-S, D.J.S., S.E.G., R.T.B., J.W., W.T., C.J.D., J.F., A.R.R., A.R.S.; Writing – Review and Editing, U.P., E.N.A., J.W., C.J.W., C.C.W., D.S., A.R.R., A.R.S.; Supervision, U.P., C.J.D., D.S., A.R.R., A.R.S.; Funding Acquisition, C.B-S., D.J.S., D.S., A.R.R., A.R.S.

## Supplemental Video Legends

Video S1. Liquid-like behavior of WNK1 puncta.

Time-lapse movie of WNK1-eGFP expressing HEK-293 cells subjected to mild hyperosmotic stress (37mM sorbitol; 337mOsm). A large spherical droplet that forms via liquid-like fusion, nucleation, and growth post-stress exhibits rapid fluorescence recovery after photobleaching (FRAP).

Video S2. WNK1 nucleation and spinodal decomposition.

Live cell imaging of two cells expressing moderate and very high levels of WNK1-eGFP, subjected to 50mM KCl, demonstrating nucleation and spinodal decomposition, respectively

Video S3. WNK1 puncta formation in mNG-WNK1 knock-in cells.

Live cell imaging of gene-edited cells expressing endogenously mNG-tagged WNK1 (Fig S2).

Video S4. Light-inducible phase separation of the extreme WNK1 C-terminus.

Live cell imaging of HEK-293 cells transfected with WNK1 constructs fused to the WT PHR domain of Cry2, and exposed to 450nm (blue) light.

Video S5. Response of CTD truncation mutants to hyperosmotic stress.

Live cell imaging of WNK1 constructs (Fig 3F), expressed in HEK-293 cells and subjected to hyperosmotic stress (50mM sorbitol).

Video S6. Hypertonic stress induces dmWNK condensate formation in *Drosophila* S2 cells.

Live cell imaging of full-length dmWNK 1-2253 (Fig 5C), expressed in S2-GAL4 cells that were initially exposed to hypotonicity (174 mOsm) and then subjected to further hypotonicity (87 mOsm, to completely dissolve condensates) or increased osmolality (125mM sorbitol, final osmolality 232 mOsm).

---

**Figure S1.**
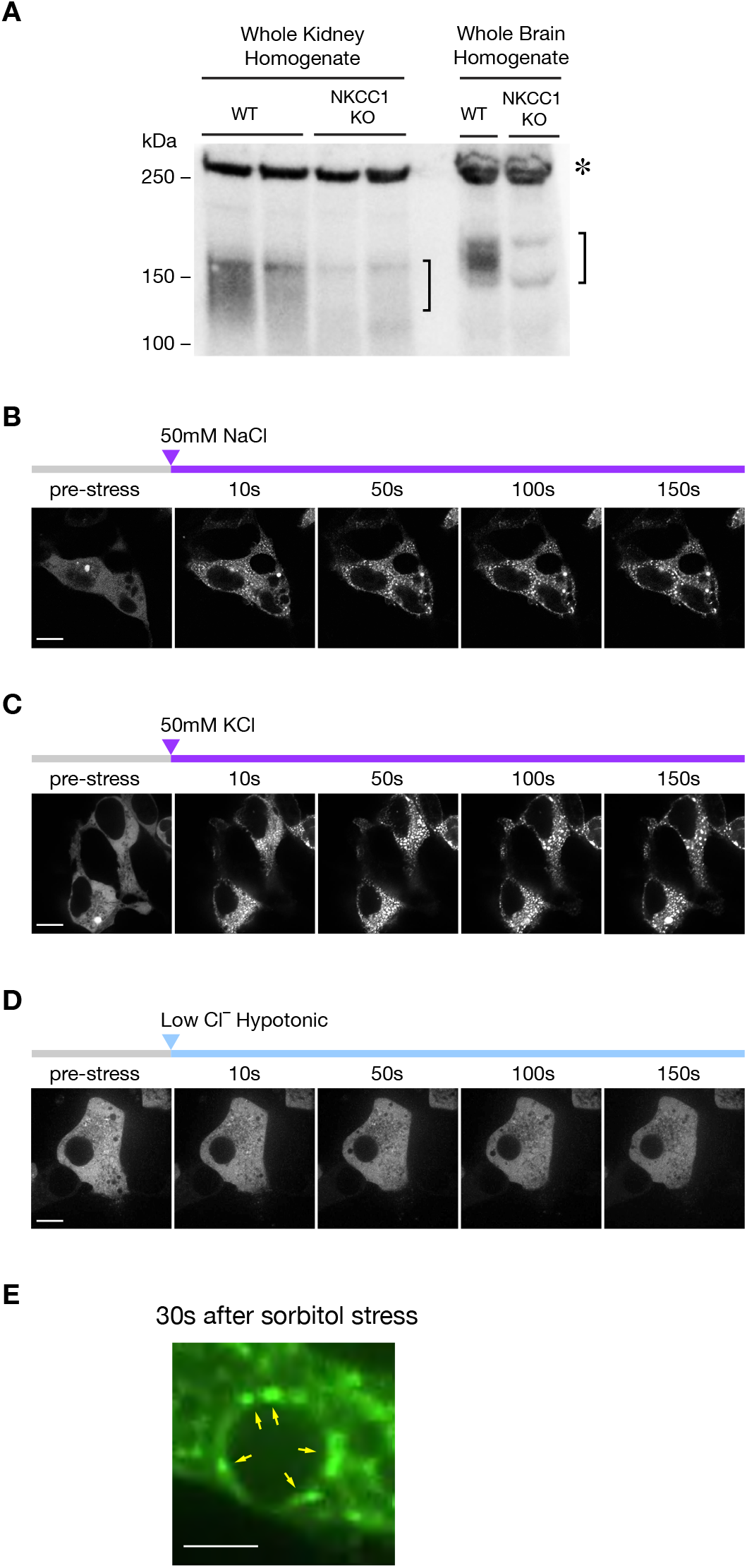
Effects of various osmotic stressors on WNK1 condensate formation. (A) Total NKCC1 antibody validation in whole kidney and brain homogenates from WT and global NKCC1 KO mice. The NKCC1 glycoprotein migrates as a broad immunodetectable band that runs slightly slower in brain tissue, as indicated with brackets. The asterisk indicates a nonspecific band. (B-C) 50mM NaCl and KCl (added to isotonic media; final osmolality = 400mOsm) both trigger the formation of WNK1 puncta. (D) No effect of low chloride hypotonic stress (isotonic media diluted 50:50 with distilled water; final osmolality = 150mOsm) on WNK1 puncta formation. (E) Consistent with liquid phase condensation, WNK1 puncta wet against intracellular surfaces (yellow arrows). Still image from Video S1. Bar = 2µm.

**Figure S2.**
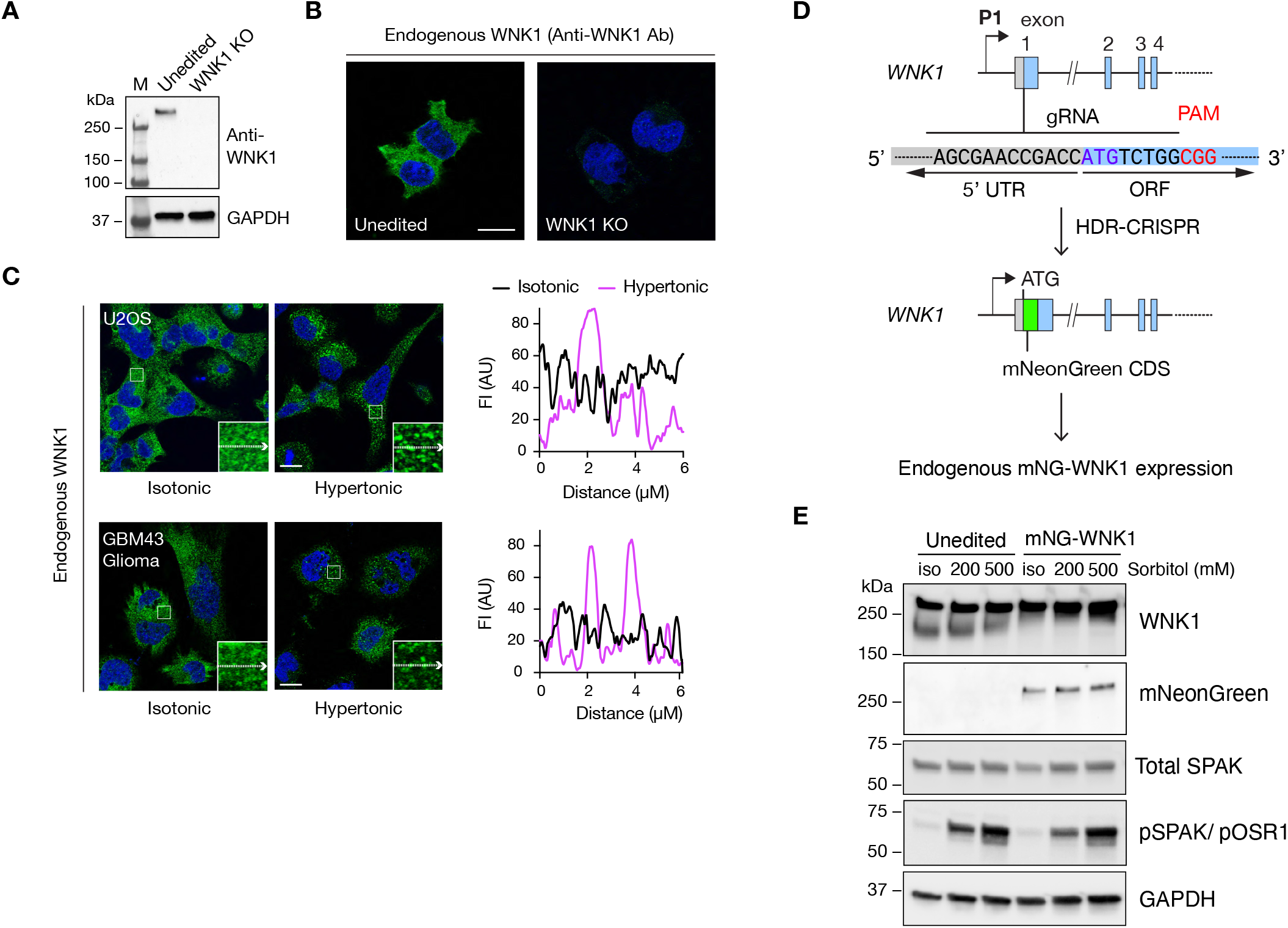
Endogenous WNK1 forms biomolecular condensates. (A-B) Validation of WNK1 antibody (RRID: AB_2679690) by immunoblotting and IF in unedited and WNK1 KO HEK cells (Roy et al., 2015b). (C) Representative fixed IF images of WNK1 puncta formation in U2OS and GBM43 cells, subjected to hypertonic stress (500mM sorbitol x 5min). Line scans correspond to fluorescence intensity (FI) traces along the trajectories of the arrows in the insets under isotonic (black) and hypertonic (magenta) conditions. Bar = 10µm. (D) CRISPR guide RNA (gRNA) sequence and exon 1 target site used to knock in an in-frame N-terminal mNeonGreen (mNG) coding DNA sequence (CDS) into the endogenous WNK1 gene locus in HEK cells. (E) Immunoblots of gene edited mNG-WNK1 cells and unedited HEK controls subjected to hypertonic stress (200 or 500mM sorbitol x 5min).

**Figure S3.**
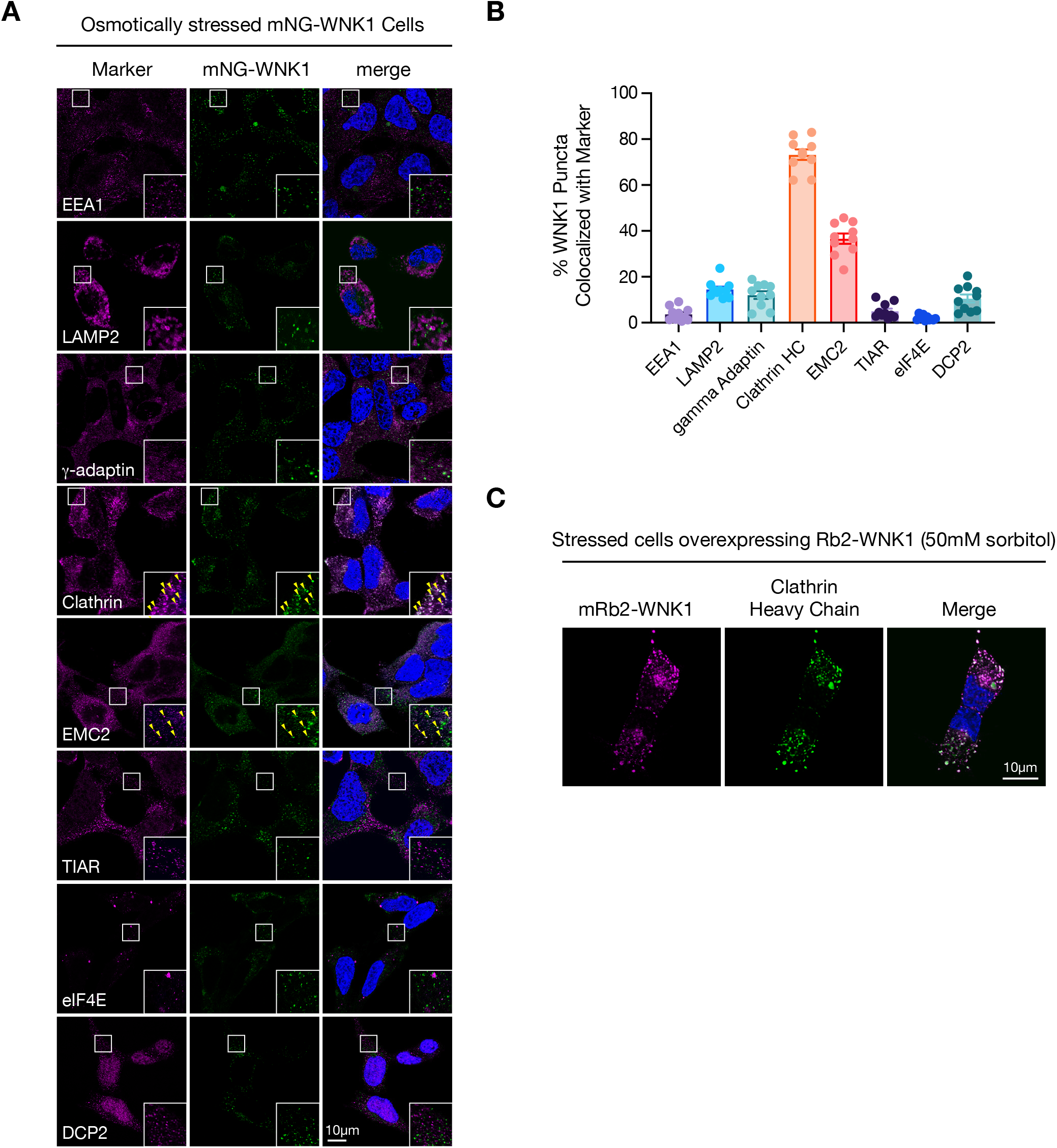
WNK1 colocalization with intracellular markers. (A) IF images of fixed HEK cells expressing endogenous mNG-tagged WNK1, subjected to hyperosmotic stress (500mM sorbitol x 5min), stained for various endogenous markers. Images were acquired with a Leica Stellaris microscope with Lightning adaptive deconvolution. (B) Quantification of colocalization (% mNG signal overlap with marker), by Mander’s M1 overlap coefficient, calculated using the JACoP plugin in FIJI. N=10 analyzed confocal fields per condition. (C) Representative Lightning deconvolution image of overexpressed mRb2-WNK1, colocalized with endogenous clathrin heavy chain in hypertonically stressed HEK cells.

**Figure S4.**
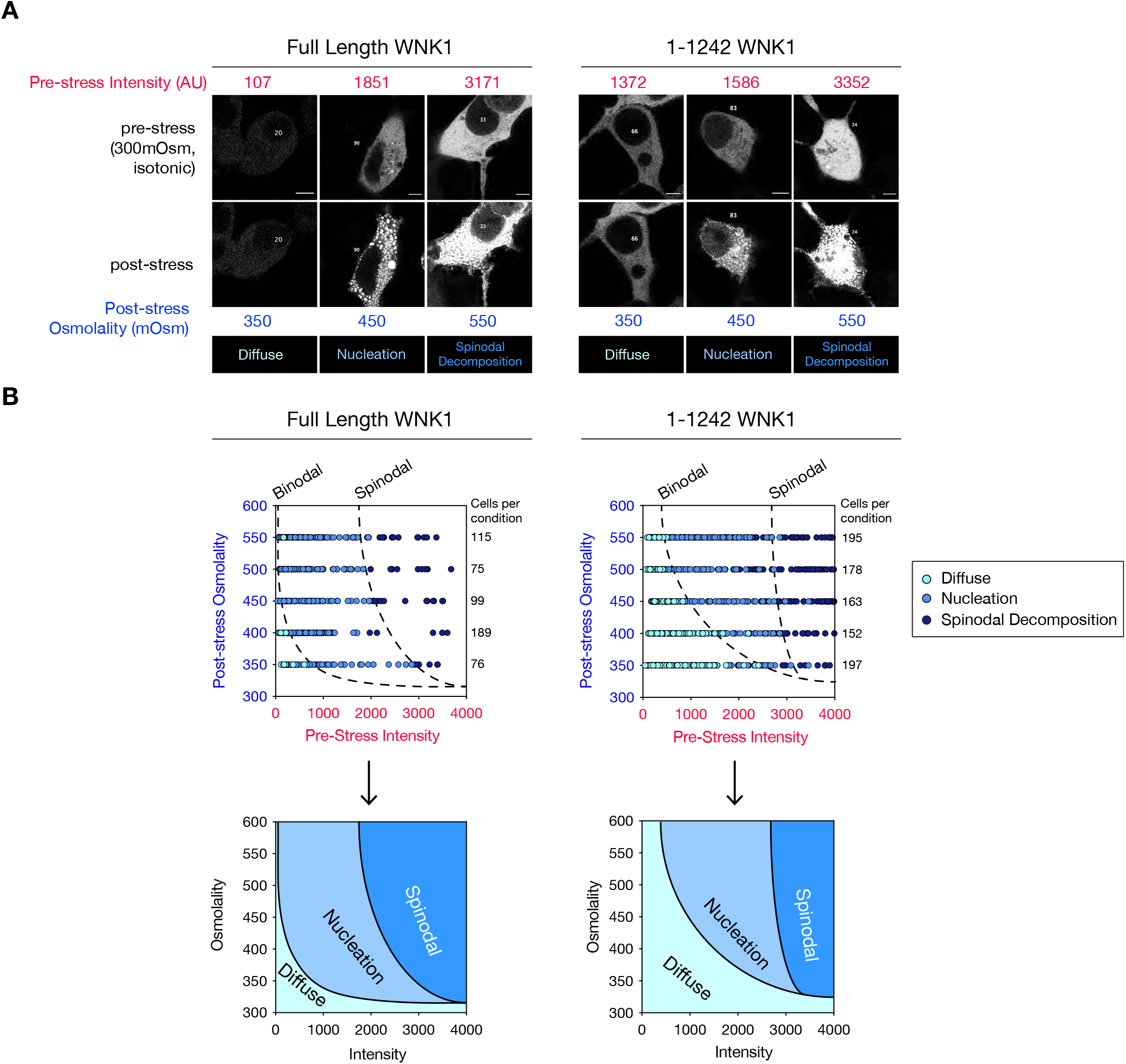
In-cell phase diagram construction. (A) Representative pre- and post-stress live cell data obtained from cells expressing full-length WNK1, or 1-1242 WNK1. Constructs were scored by a panel of blinded observers as undergoing no phase separation (diffuse), nucleation, or spinodal decomposition, depending on morphology, as shown. Individual cells were assigned numbers for the purposes of deidentification and blinded scoring, as described in the Methods. Pre-stress fluorescence intensities (red) were plotted relative to osmotic stress (blue) to generate phase diagrams. (B) Raw phase diagram data, as plotted for full length WNK1 and for 1-1242 WNK1. Binodal and spinodal phase boundaries were drawn at fluorescence intensities where the WNK1 constructs exclusively formed nucleated droplets, or underwent spinodal decomposition, respectively.

**Figure S5.**
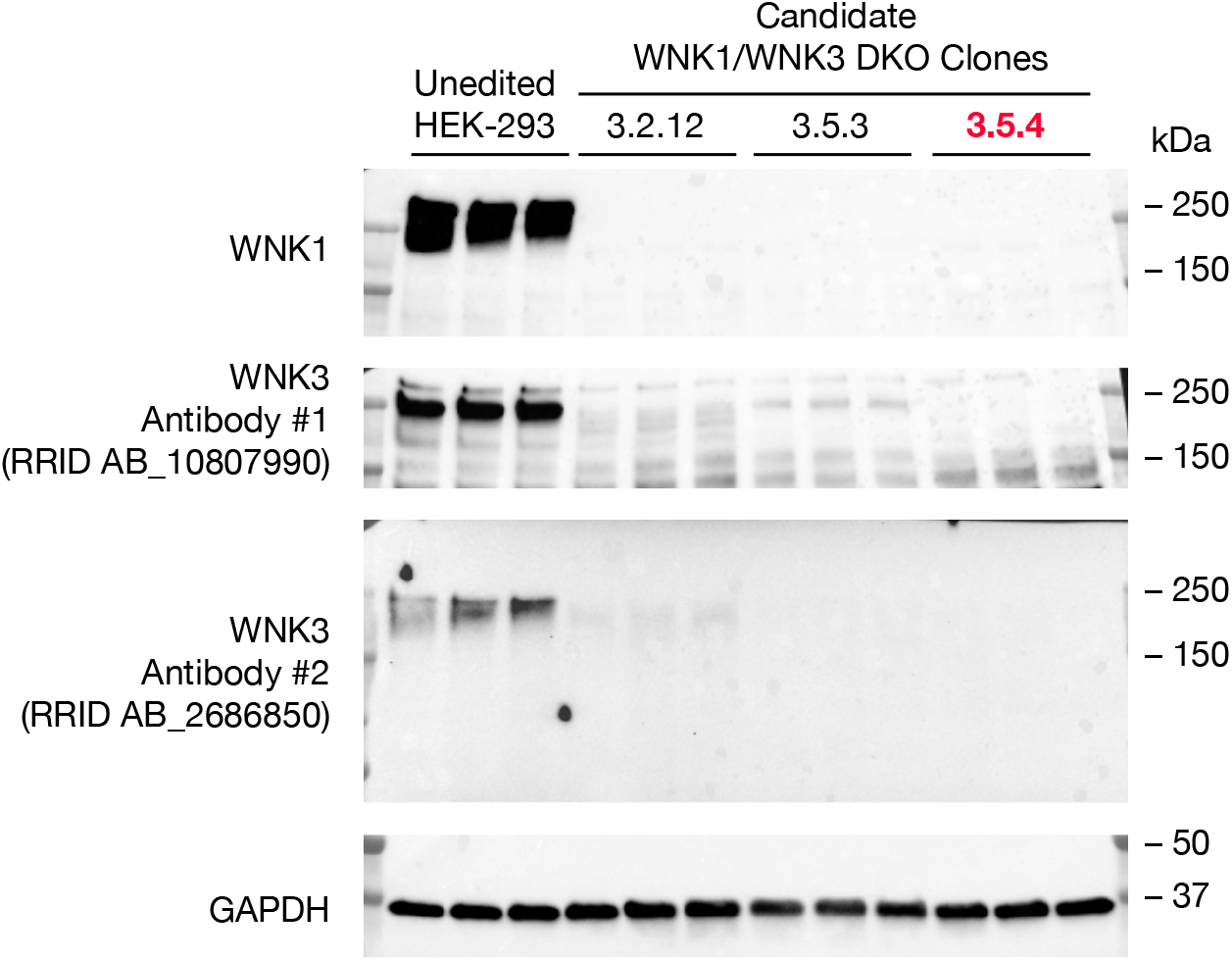
WNK1/WNK3 Double knockout cells. Immunoblots of candidate WNK1/WNK3 DKO HEK cell clones, compared to unedited HEK-293 cell controls. Clone 3.5.4 (red) was used for this study.

**Figure S6.**
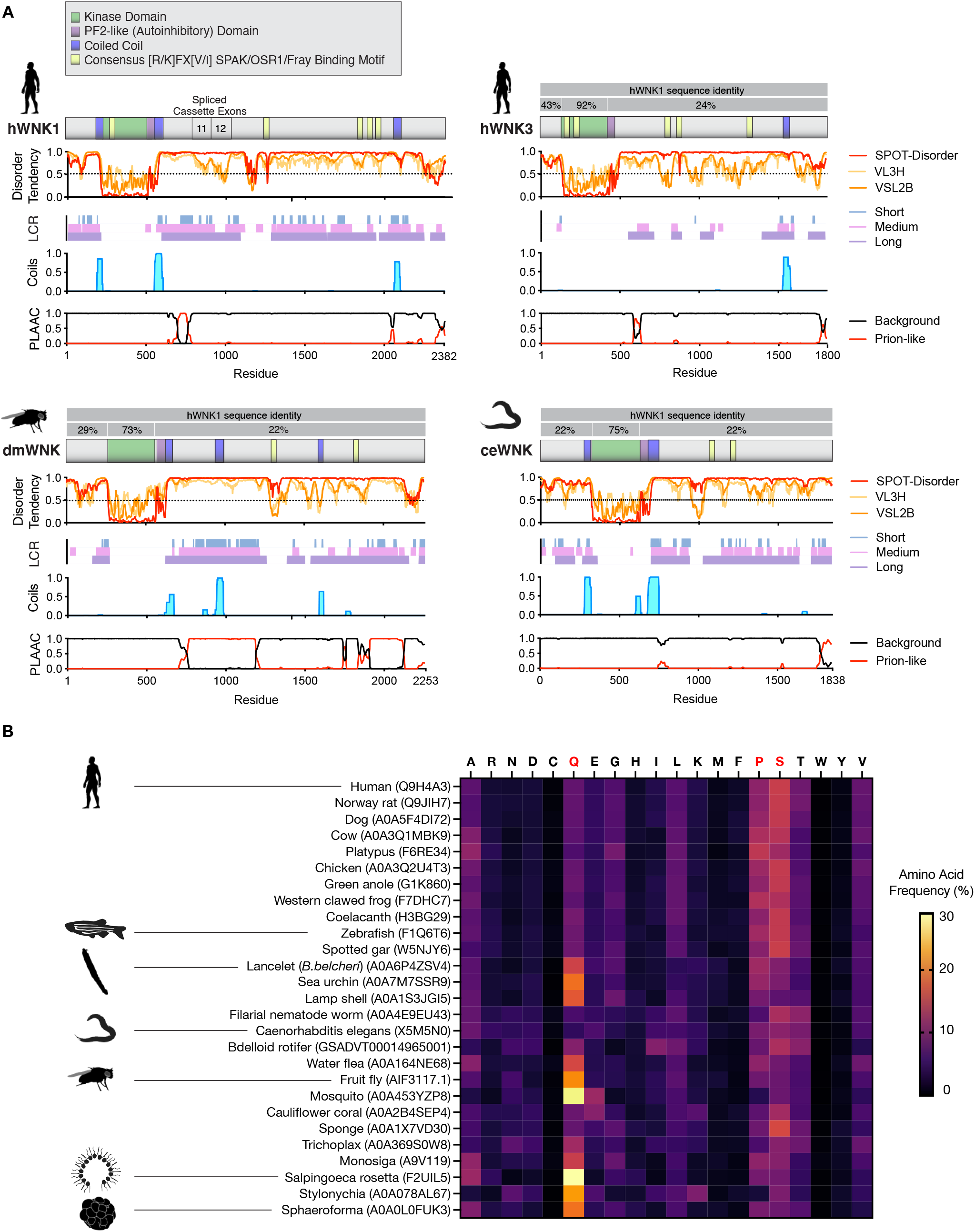
Evolutionary analysis of the WNK1 CTD. (A) Disorder, LCR, Coiled coil, and PLD analysis of human WNK1 (hWNK1), human WNK3 (hWNK3), *Drosophila melanogaster* WNK (dmWNK), and *Caenorhabditis elegans* WNK (ceWNK). Sequence identity of hWNK3, dmWNK, and ceWNK relative to hWNK1, divided into three regions: the N-terminus upstream of the kinase domain, the kinase domain itself, and the large C-terminal domain (CTD) downstream from the kinase domain. Despite the poor sequence identity of the CTDs, the regional disorder tendencies and PLDs are nearly identical. (B) Amino acid composition of WNK CTDs across evolution, ranging from humans to protists. In higher organisms that harbor multiple WNK paralogs, WNK1 was used for this analysis. Note that the CTD composition in higher organisms is proline/serine biased, whereas in organisms predating vertebrate animals, the CTD is frequently highly enriched in glutamine. Most accession numbers are UniProt identifiers, with exceptions, as noted in Table S1.

**Table S1.** Cross-species identification of WNK kinases. WNK kinases were defined as having an atypically placed catalytic lysine (K233 in human) and structurally-defined hydrophobic residues in the kinase domain that form a chloride binding pocket (F283/ L299/ L369/ L371 in human).

| Common | Taxonomic | Source | Accession/ID | Name | Protein Length (AA) | Aligned Catalytic Lysine | Aligned CI- binding residues | CTD boundary† |
| --- | --- | --- | --- | --- | --- | --- | --- | --- |
| Human | *Homo sapiens* | UniProtKb (Swiss-Prot) | Q9H4A3 | WNK1 | 2382 | K233 | F283/L299/L369/L371 | 556-end |
| Rat | *Rattus norvegicus* | UniProtKb (Swiss-Prot) | Q9JIH7 | WNK1 | 2126 | K233* | F283/L299/L369/L371* | 556-end |
| Dog | *Canis familiaris* | UniProtKb (TrEMBL) | A0A5F4DI72 | WNK1 | 2847 | K238 | F288/L304/L374/L376 | 561-end |
| Cow | *Bos taurus* | UniProtKb (TrEMBL) | A0A3Q1MBK9 | WNK1 | 2832 | K238 | F288/L304/L374/L376 | 561-end |
| Platypus | *Ornithorhynchus anatinus* | UniProtKb (TrEMBL) | F6RE34 | WNK1 | 2228 | K169 | F219/L235/L305/L307 | 492-end |
| Chicken | *Gallus gallus* | UniProtKb (TrEMBL) | A0A3Q2U4T3 | WNK1 | 2662 | K238 | F288/L304/L374/L376 | 561-end |
| Green anole | *Anolis carolinensis* | UniProtKb (TrEMBL) | G1K860 | WNK1 | 2864 | K207 | F257/L273/L343/L345 | 530-end |
| Western clawed frog | *Xenopus tropicalis* | UniProtKb (TrEMBL) | F7DHC7 | WNK1 | 2272 | K196 | F246/L262/L332/L334 | 519-end |
| Coelacanth | *Latimeria chalumnae* | UniProtKb (TrEMBL) | H3BG29 | WNK1 | 2654 | # | F31/L47/L117/L119 | 304-end |
| Zebrafish | *Danio rerio* | UniProtKb (TrEMBL) | F1Q6T6 | wnk1b | 2357 | K226 | F276/L292/L362/L364 | 549-end |
| Spotted gar | *Lepisosteus oculatus* | UniProtKb (TrEMBL) | W5NJY6 | Non-specific serine/threonine protein kinase | 2090 | K230 | F280/L296/L366/L368 | 553-end |
| Lancelet | *Branchiostoma belcheri* | UniProtKb (TrEMBL) | A0A6P4ZSV4 | Non-specific serine/threonine protein kinase | 2587 | K228 | F278/L294/L364/L366 | 552-end |
| Sea urchin | *Strongylocentrotus purpuratus* | UniProtKb (TrEMBL) | A0A7M7SSR9 | Non-specific serine/threonine protein kinase | 2622 | K260 | F310/L327/L397/L399 | 581-end |
| Lamp shell | *Lingula unguis* | UniProtKb (TrEMBL) | A0A1S3JGI5 | Non-specific serine/threonine protein kinase | 2996 | K368 | F418/M435/L505/L507 | 696-end |
| Filarial nematode | *Brugia malayi* | UniProtKb (TrEMBL) | A0A4E9EU43 | Bma-wnk-1 | 1741 | K210 | F266/L284/L354/L356 | 549-end |
| Roundworm | *Caenorhabditis elegans* | UniProtKb (Swiss-Prot) | X5M5N0 | WNK1 | 1850 | K346 | F396/L414/L484/L486 | 679-end |
| Rotifer | *Adineta vaga* | Ensembl Metazoa | GSADVT00014965001 | – | 1323 | K449 | F500/L518/L592/L594 | 754-end |
| Water flea | *Daphnia magna* | UniProtKb (TrEMBL) | A0A164NE68 | Non-specific serine/threonine protein kinase | 2540 | K364 | F547/L563/L635/L635 | 824-end |
| Fruit fly | *Drosophila melanogaster* | NCBI | AIF73117.1 | Wnk variant 1 | 2253 | K285 | F335/L351/L421/L423 | 612-end |
| Mosquito | *Anopheles gambiae* | UniProtKb (TrEMBL) | A0A453YZP8 | Non-specific serine/threonine protein kinase | 3443 | K562 | F612/L631/L701/L703 | 892-end |
| Cauliflower coral | *Stylophora pistillata* | UniProtKb (TrEMBL) | A0A2B4SEP4 | WNK1 | 1663 | K91 | F140/L158/L228/L230 | 422-end |
| Sponge | *Amphimedon queenslandica* | UniProtKb (TrEMBL) | A0A1X7VD30 | Non-specific serine/threonine protein kinase | 1703 | K169 | L219/L237/L307/L309 | 492-end |
| – (Placozoan) | *Trichoplax adhaerens* | UniProtKb (TrEMBL) | A0A369S0W8 | Non-specific serine/threonine protein kinase | 1565 | K132 | F182/L203/L275/L277 | 454-end |
| – (Choanoflagellate) | *Monosiga brevicollis* | UniProtKb (TrEMBL) | A9V119 | Predicted protein | 1239 | K171 | F223/F236/L307/L309 | 503-end |
| – (Choanoflagellate) | *Salpingoeca rosetta* | UniProtKb (TrEMBL) | F2UIL5 | WNK protein kinase | 1767 | K486 | Y534/V550/M623/L625 | 831-end |
| Ciliate | *Stylonychia lemnae* | UniProtKb (TrEMBL) | A0A078AL67 | Mitogen activated protein kinase family | 1891 | K281 | F331/F345/L416/L418 | 575-end |
| – (Ichthyosporean) | *Sphaeroforma arctica* | UniProtKb (TrEMBL) | A0A0L0FUK3 | WNK protein kinase | 1295 | K8 | C58/F81/L151/L153 | 262-end |

\*Rat WNK1 was used as the template for amino acid alignments.
Conservative and nonconservative substitutions are noted in blue and red, respectively.
†CTD boundaries = the first residue C-terminal to the PF2-like domain (residue 556 in human) extending to the C-terminal end of the protein.

**Table S2.**
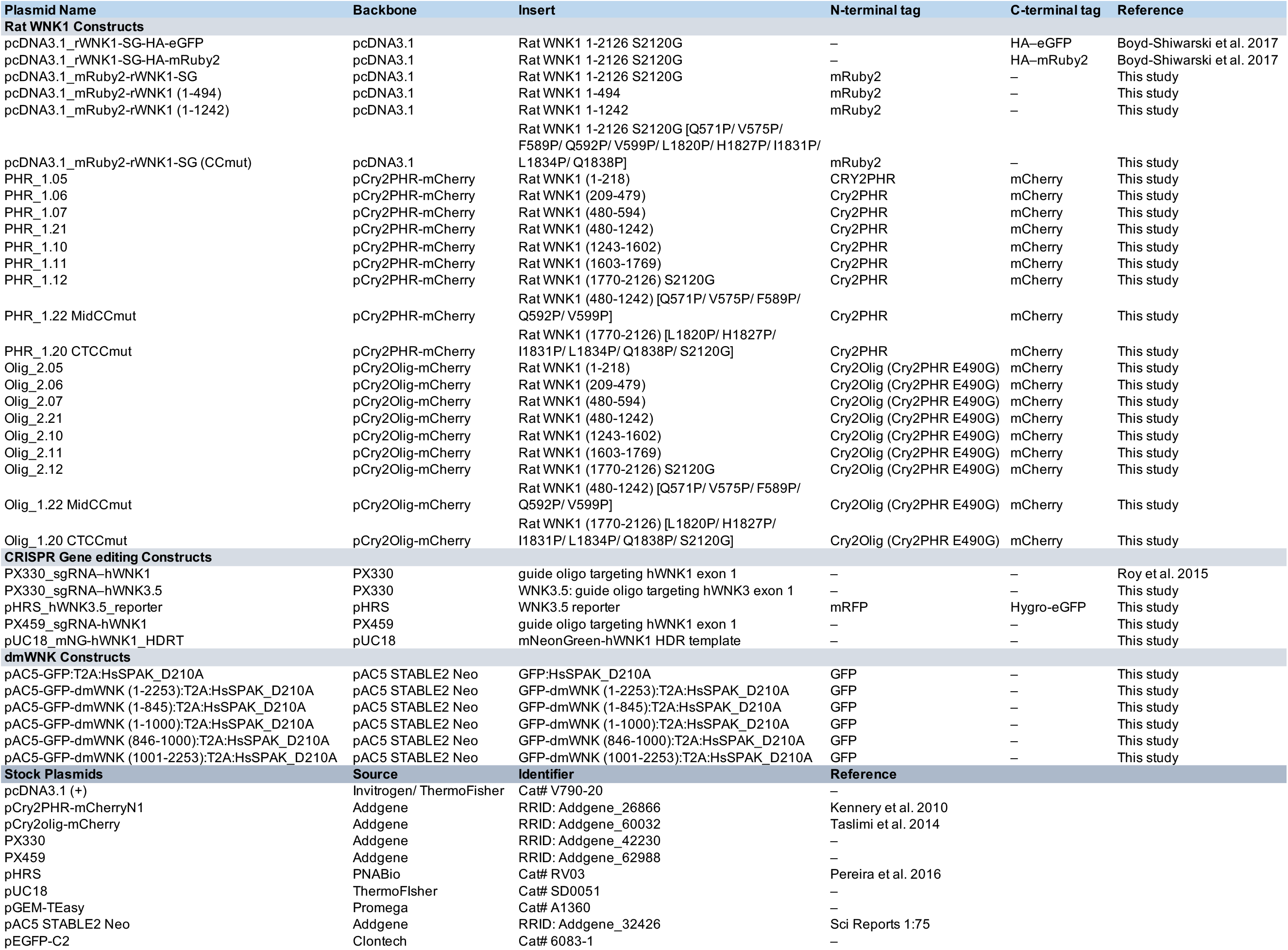
Plasmids.

**Table S3.** Antibodies.

| Antibody Name | Host | Source | Cat # | RRID | Dilution (WB) | Dilution (IF) |
| --- | --- | --- | --- | --- | --- | --- |
| **Primary Antibodies** |  |  |  |  |  |  |
| WNK1 | Rabbit | Atlas | HPA059157 | AB_2683935 | 1:2000 | 1:200 |
| WNK1 | Rabbit | Atlas | HPA046541 | AB_2679690 | 1:1000 | 1:200 |
| WNK3 | Rabbit | Millipore Sigma | 07-2262 | AB_10807990 | 1:1000 | – |
| WNK3 | Rabbit | Millipore Sigma | HPA077678 | AB_2686850 | 1:1000 | 1:200 |
| OSR1 | Rabbit | Atlas | HPA008237 | AB_1079540 | – | 1:200 |
| SPAK | Rabbit | Cell Signaling Technology | 2281 | AB_2196951 | 1:1000 | 1:200 |
| phospho-SPAK (Ser373) / phospho-OSR1 (Ser325) | Rabbit | Millipore Sigma | 07-2273 | AB_11205577 | 1:1000 | 1:200 |
| NKCC1 | Guinea pig | (This paper) | – | – | 1:2000 | – |
| pNCC (Thr58)* (Crossreacts with pNKCC1) | Rabbit | Jan Loffing (Sorensen et al 2013) | – | – | 1:4000 | – |
| pKCC (Thr1007) | Rabbit | PhosphoSolutions | p1551-1007 | AB_2716769 | 1:1500 | – |
| mNeonGreen [32F6] | Mouse | Bulldog Bio | MNG32F6 | AB_2827566 | 1:2000 | – |
| tRFP | Rabbit | Evrogen | AB233 | AB_2571743 | 1:3000 | – |
| GAPDH | Rabbit | Cell Signaling Technology | 2118 | AB_561053 | 1:10,000 | – |
| EEA1 | Mouse | BD Biosciences | 610456 | AB_397829 | – | 1:200 |
| LAMP2 | Mouse | DS Hybridoma Bank | H4B4 | AB_528129 | – | 1:200 |
| gamma-adaptin (AP-1) | Mouse | BD Transduction Lab | 610385 | AB_397768 | – | 1:200 |
| Clathrin Heavy Chain | Rabbit | Abcam | AB21679 | AB_2083165 | – | 1:200 |
| EMC2 | Rabbit | Proteintech | 25443-1-AP | AB_2750836 | – | 1:200 |
| TIAR | Mouse | (Daigle et al., 2016) | – | – | – | 1:200 |
| eIF4e | Rabbit | Bethyl Laboratories | IHC-00127-T | AB_2246256 | – | 1:200 |
| DCP2 | Rabbit | Bethyl Laboratories | A302-597A-T | AB_10555903 | – | 1:200 |
| **Secondary Antibodies** |  |  |  |  |  |  |
| Peroxidase AffiniPure Goat Anti-Rabbit IgG (H+L) | Goat | Jackson ImmunoResearch Laborato | 111-035-003 | AB_2313567 | 1:5000 | – |
| Peroxidase AffiniPure Goat Anti-Guinea Pig IgG (H+L) | Goat | Jackson ImmunoResearch Laborato | 106-035-003 | AB_2337402 | 1:5000 | – |
| Peroxidase AffiniPure Goat Anti-Mouse IgG (H+L) | Goat | Jackson ImmunoResearch Laborato | 115-035-003 | AB_10015289 | 1:5000 | – |
| Goat anti Rabbit Alexa 488 | Goat | Invitrogen | A11008 | AB_143165 | – | 1:200 |
| Donkey anti Rabbit Cy3 | Donkey | Biolegend | 406402 | AB_893532 | – | 1:200 |
| Goat anti Mouse Alexa 568 | Goat | Invitrogen | A11004 | AB_2534072 | – | 1:200 |

